# VIP-producing enteric neurons interact with innate lymphoid cells to regulate feeding-dependent intestinal epithelial barrier functions

**DOI:** 10.1101/721464

**Authors:** Jhimmy Talbot, Paul Hahn, Lina Kroehling, Henry Nguyen, Dayi Li, Dan R. Littman

## Abstract

The intestinal mucosa serves as both a conduit for uptake of food-derived nutrients and microbiome-derived metabolites and as a barrier that prevents tissue invasion by microbes and tempers inflammatory responses to the myriad contents of the lumen. How the intestine coordinates physiological and immune responses to food consumption to optimize nutrient uptake while maintaining barrier functions remains unclear. Here, we describe how a gut neuronal signal triggered by food intake is integrated with intestinal antimicrobial and metabolic responses controlled by type 3 innate lymphoid cells (ILC3)^1-3^. Food consumption rapidly activates a population of enteric neurons that express vasoactive intestinal peptide (VIP)^4^. Projections of VIP-producing neurons (VIPergic neurons) in the lamina propria are in close proximity to clusters of ILC3 that selectively express VIP receptor type 2 (VIPR2 or VPAC2). ILC3 production of IL-22, which is up-regulated by commensal microbes such as segmented filamentous bacteria (SFB)^5-7^, is inhibited upon engagement of VIPR2. As a consequence, there is a reduction in epithelial cell-derived antimicrobial peptide, but enhanced expression of lipid-binding proteins and transporters^8^. During food consumption, activation of VIPergic neurons thus enhances growth of epithelial-associated SFB and increases lipid absorption. Our results reveal a feeding- and circadian-regulated dynamic intestinal neuro-immune circuit that promotes a trade-off between IL-22-mediated innate immune protection and efficiency of nutrient absorption. Modulation of this pathway may hence be effective for enhancing resistance to enteropathogen^2,3,9^ and for treatment of metabolic diseases.

## MAIN TEXT

Type 3 innate lymphoid cells (ILC3) promote maintenance of intestinal immune and metabolic homeostasis by integrating cytokine-mediated and glia-derived cues and, through the production of IL-22 and other cytokines, conveying information from the luminal microbiota to intestinal epithelial cells (IECs) and cells in the lamina propria ^1-3,5,10^. The activation of ILC3 is regulated by cytokines, including IL-23, IL-1β, and TL1A, which are produced by mononuclear phagocytes and induced by intestinal microbe-derived stimuli^5,7,11^. Cytokines produced by activated ILC3, particularly IL-22, support the production of antimicrobial peptides (*e.g. RegIIIγ*) and mucin by IECs, ensuring spatial segregation of microbes from the intestinal tissue^12,13^. This microbiota-ILC3-IEC circuit promotes intestinal barrier function by controlling intestinal commensal microbiota and mediating rapid protective responses to enteropathogens^2,3,9,12^. Different subtypes of ILC3s are present within the intestinal lamina propria and are dispersed or in tertiary lymphoid tissue clusters known as cryptopatches (CPs) and isolated lymphoid follicles (ILFs)^14,15^. In perusing transcriptomic datasets of small intestinal ILCs, we noticed in CCR6^+^ ILC3, which comprise the lymphoid tissue inducer (LTi) cells enriched in CPs and ILFs, selective expression of multiple neurotransmitter/neuropeptide receptors and genes related to axonal guidance and neuron differentiation when compared to CCR6^neg^ ILC3 and NCR^+^ ILC3 (Fig. 1a and Extended Data Fig. 1a-c)^16,17^. This prompted us to evaluate whether CCR6^+^ ILC3 in the intestinal lamina propria were associated with neuronal projections from the enteric nervous system (ENS). The ENS is part of the peripheral autonomic nervous system, which controls gastrointestinal functions by promoting rapid gut responses to changes in the luminal compartment (e.g. food intake, microbes, metabolites, etc.^18-22^). We observed that ILC3 in CPs/ILFs were in close proximity to neuronal projections in the lamina propria (Fig. 1b-d and Supplementary Videos 1,2). The CP/ILF-associated neuronal projections were positive for vasoactive intestinal peptide upon staining with specific antibody (VIP, Fig. 1e and Extended Data Fig. 2). We also observed association of VIPergic neurons and colonic lymphoid patches (Extended Data Fig. 2 a,b). Indeed, ILC3 in CPs/ILFs were closer to VIPergic neurons than to neurons positive for substance P or tyrosine hydroxylase (adrenergic) (Fig. 1f and Extended Data Fig. 3). These anatomical findings, together with the observation that CCR6^+^ ILC3, but not intestinal Th17 or γδT17 cells, express a high amount of VIP receptor type 2 (*Vipr2*) (Fig. 1a and extended Data Figure 4a), prompted us to investigate whether signaling from VIPergic neurons could modulate CCR6^+^ ILC3 functions and intestinal immune homeostasis.

**Figure 1.**
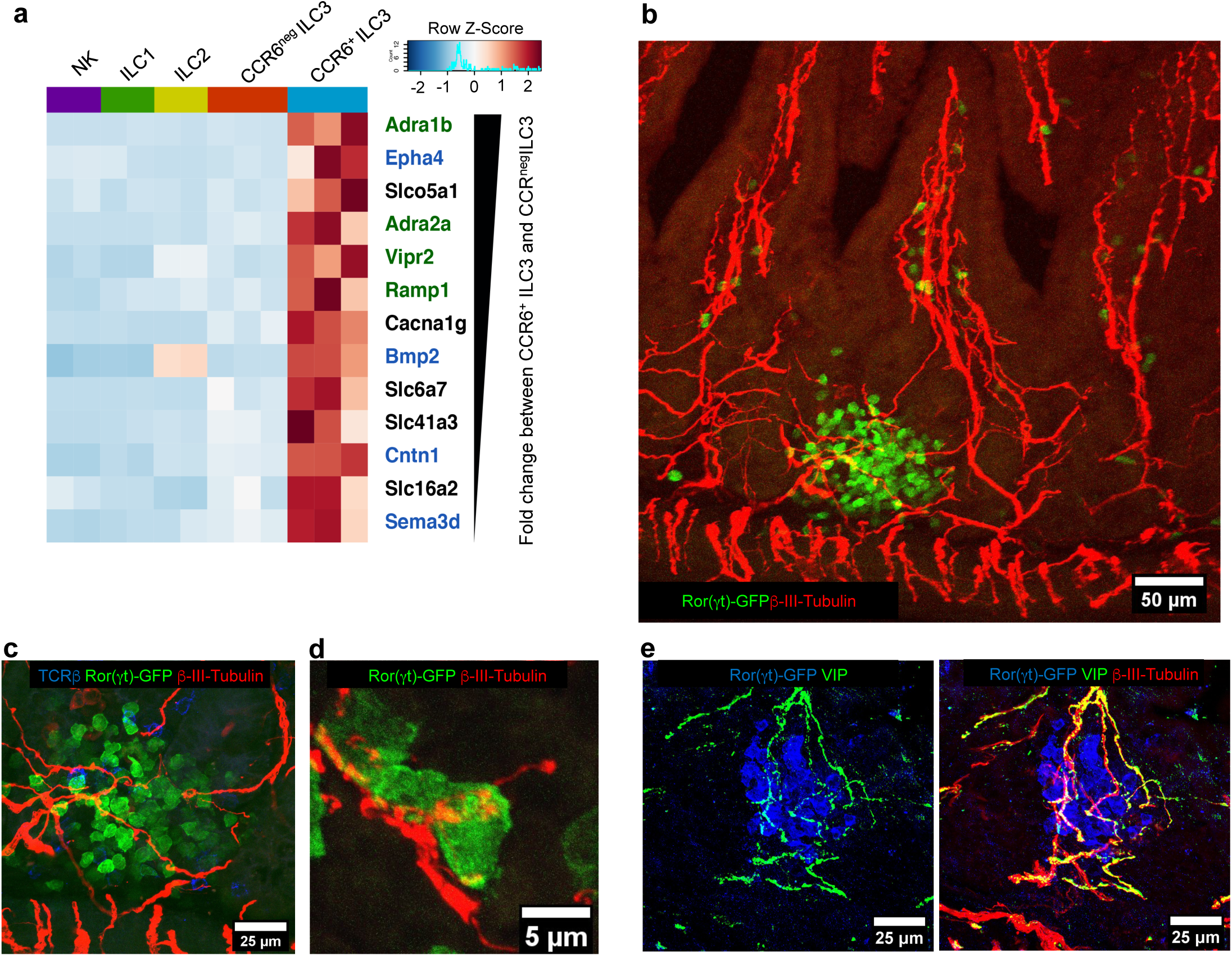
Processes of VIP-producing enteric neurons are in close proximity to *Vipr2*-expressing ILC3 within cryptopatches. **a**, Heatmap of differentially expressed neural-related genes between intestinal CCR6^+^ ILC3 and CCR6^neg^ ILC3 (Fold change ≥ 2, *p-value <0.05*, GSE116092). Color scale is based on normalized read counts. Genes are listed on the right hand margin ranked based on the relative fold change, and color coded: Green: Neurotransmitter/neuropeptide receptors, Blue: genes related to nervous system development/axonal guidance and contact. Expression in other ILC subsets is included for comparison. **b-d**, Representative confocal images from the small intestine of *RORc(t)*^EGFP/+^ mice show clusters of intestinal ILC3 (cryptopatch) in close proximity to enteric neurons in the small intestine lamina propria (see supplementary videos 1 and 2). Pan-neuronal marker: β3-tubulin^+^ (red), ILC3: GFP^+^TCRβ^neg^ (green and blue, respectively). n=4 mice, all the 110 ILC3 clusters analyzed exhibited β3-tubulin^+^ fibers interacting with ILC3. **e**, Neurochemical code of cryptopatch-associated enteric neurons. Representative confocal images from the small intestine of *RORc(t)*^EGFP/+^ mice show cryptopatch-associated enteric neurons are positive for VIP. Neurons: β3-tubulin^+^ (red), VIP^+^ (green); ILC3: GFP^+^ (blue). n=4 mice, 40 ILC3 clusters analyzed, with all displaying VIP^+^ β3-tubulin^+^ fibers interacting with ILC3.

We isolated ILC3 from the lamina propria of the small intestine of C57BL/6 mice (Extended Data Fig. 4b) and observed that *in vitro* activation of VIPR2 inhibited IL-23-induced production of IL-22 and IL-17A by CCR6^+^ ILC3 (Fig. 2a,b and Extended Data Fig. 4c-e), although there was no effect on cell activation (marked by Sca-1 up-regulation^5^) or on IL-22 production by CCR6^neg^ ILC3 (Extended Data Fig. 4f-h). In order to investigate the *in vivo* role of VIPR2 activation on IL-22 production in unstimulated (4h/37C) CCR6^+^ ILC3 functions, we generated mixed bone marrow chimeric mice reconstituted with a 1:1 ratio of isotype-marked *Vipr2*^+/+^ and *Vipr2*^-/-^ cells. IL-22 production (assessed following inhibition of Golgi trafficking in unstimulated *ex vivo* cultured cells) was observed in a greater proportion of *Vipr2*^-/-^ CCR6^+^ ILC3 when compared to *Vipr2*^+/+^ CCR6^+^ ILC3 (Extended Data Fig 5a,b). We evaluated the transcriptional profile of *Vipr2*^+/+^ and *Vipr2*^-/-^ CCR6^+^ ILC3 from these mice and observed a marked difference between them, with up-regulation of genes involved in retinoic acid-mediated signaling (*Rxra*) and translation initiation (*Eif5a*) in the absence of *Vipr2* (Extended Data Fig 5c,d,e). We also generated mice with conditional deletion of the gene encoding VIPR2 in ILC3 (*RORc(t)*^*Cre*^*Vipr2*^*fl/fl*^) and found that a larger proportion and number of unstimulated CCR6^+^ ILC3 isolated from the ileum of these mice produced IL-22 than cells isolated from WT (*RORc(t)*^*Cre*^*Vipr2*^*+/+*^) littermates (Fig. 2c and Extended Data Fig. 5f-i). There was also a higher frequency of cells positive for IL-17A in CCR6^+^ ILC3 from *RORc(t)*^*Cre*^*Vipr2*^*fl/fl*^ mice, although the amount was low compared to IL-22 (Extended Data Fig. 5j,k). No differences were observed in the frequency of IL-17A or IL-22 production among CD3^+^RORγt^+^ cells (γδT17 and Th17 cells) (Extended Data Fig. 5l,m). *RegIIIγ* mRNA, regulated by IL-22, was considerably higher in ileal extracts enriched for intestinal epithelial cells (IECs) from *RORc(t)*^*Cre*^*Vipr2*^*fl/fl*^ mice when compared to *RORc(t)*^*Cre*^*Vipr2*^*+/+*^ mice (Fig. 2d), suggesting that VIPergic signaling through VIPR2 inhibits IL-22-mediated innate immune responses and epithelial cell reactivity. In addition, small intestine villi and crypt lengths, as well as frequency of proliferating cells (Ki67^+^) in the crypts, were increased in *RORc(t)*^*Cre*^*Vipr2*^*fl/fl*^ mice (Extended Data Fig. 5n-q), consistent with observations of the *in vivo* effects of chronic IL-22 overexpression or treatment^23,24^.

**Figure 2.**
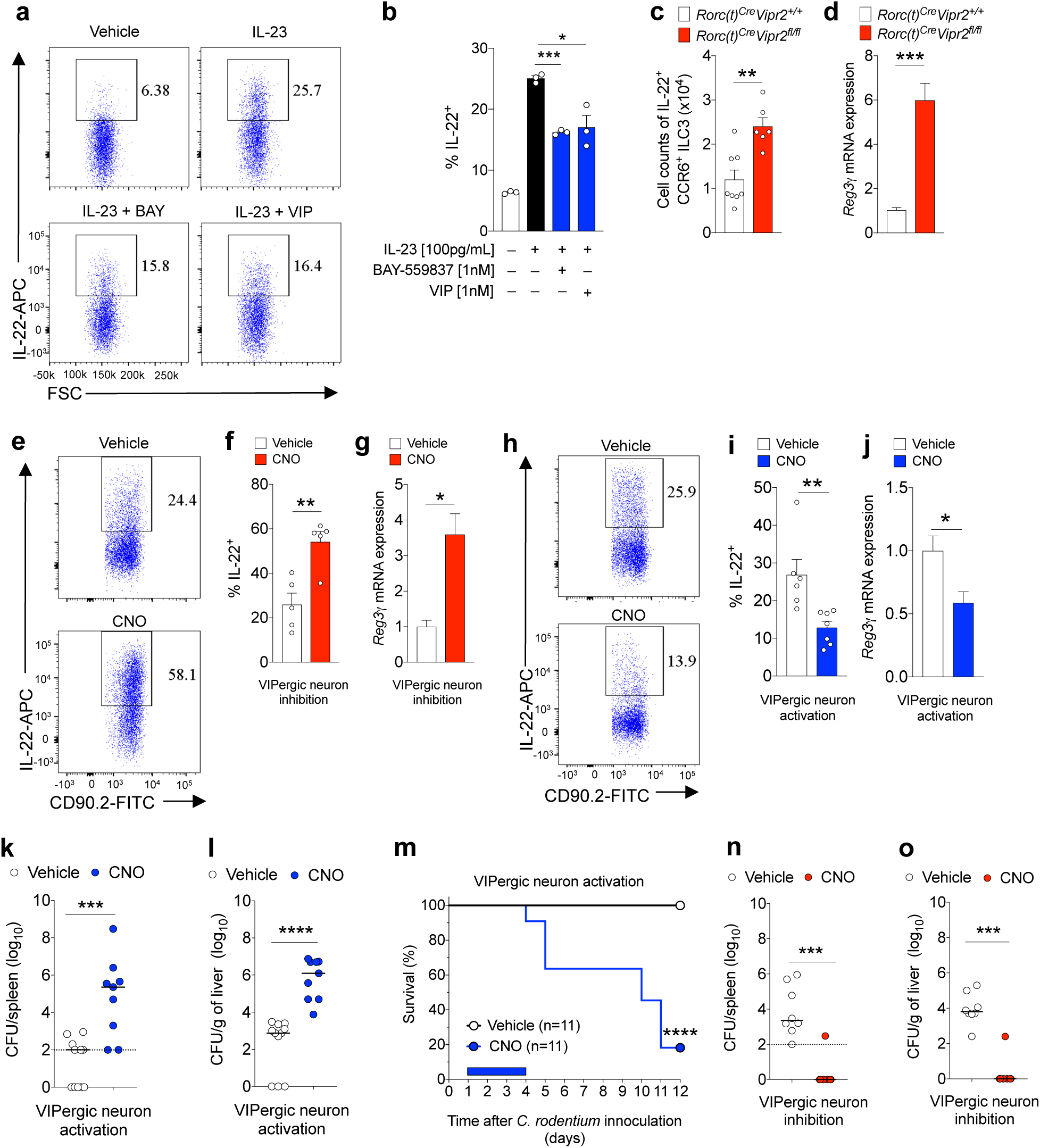
VIPergic signaling promotes reduction of mucosal barrier function by VIPR2-dependent inhibition of CCR6^+^ ILC3. **a, b**, Representative FACS plot (**a**) and summaries (**b**) indicating IL-22 expression in purified CCR6^+^ ILC3 after *in vitro* IL-23 stimulation with/without VIPR2 agonist ligands. BAY: BAY-559837, VIP: vasoactive intestinal peptide. \**P*<0.05 and \*\*\**P*<0.001 (*t-test*). Data are representative of three independent experiments. **c**, Number of IL-22^+^ CCR6^+^ ILC3 present in the ileum of *RORc(t)*^*Cre*^*Vipr2*^*+/+*^ and *RORc(t)*^*Cre*^ *Vipr2*^*fl/f*^ mice. *RORc(t)*^*Cre*^*Vipr2*^*+/+*^: 8; *RORc(t)*^*Cre*^ *Vipr2*^*fl/f*^: 6. Data are representative of 2 independent experiments, \*\**P*<0.01 (*t-test*). **d**, Normalized *RegIIIγ* mRNA expression in IEC-enriched ileal extracts from *RORc(t)*^*Cre*^*Vipr2*^*+/+*^ and *RORc(t)*^*Cre*^ *Vipr2*^*fl/f*^ mice. *RORc(t)*^*Cre*^*Vipr2*^*+/+*^: 5; *RORc(t)*^*Cre*^ *Vipr2*^*fl/f*^: 5. Data are representative of 3 independent experiments,\*\*\**P*<0.001 (*t-test*). **e-g**, Effect of VIPergic inhibition on ILC3 and IECs. Representative FACS plot (**e**) and summary (**f**) indicating IL-22 expression in CCR6^+^ ILC3 from the ileum of *Vip*^*IRES-Cre*^*hM4Di*^*fl-stop-fl/+*^ mice (DREADD for VIPergic inhibition) at 24h following treatment with vehicle or CNO (Clozapine-N-oxide, DREADD ligand). Vehicle: n=5, CNO: n=5. \*\**P*<0.01 (*t-test*). Data representative of three independent experiments. **g**, Normalized *RegIIIγ* mRNA expression in ileal extracts enriched for IECs at 24h following treatment. Vehicle: n=3, CNO: n=3. \**P*<0.05 (*t-test*). Data representative of two independent experiments. **h-j**, Effect of VIPergic activation on ILC3 and ileal extracts enriched for IECs. Representative FACS plot (**h**) and summaries (**i**) indicating IL-22 expression in CCR6^+^ ILC3 from ileum of *Vip*^*IRES-Cre*^*hM3Dq*^*fl-stop-fl/+*^ (DREADD for VIPergic activation) at 24h following the treatment with vehicle or CNO. Vehicle: 6, CNO: 7. \*\**P*<0.01 (*t-test*). **j**, Normalized *RegIIIγ* mRNA expression in ileal extracts enriched for IECs at 24h following treatment with vehicle or CNO. Vehicle: 7, CNO: 6. \**P*<0.05 (*t-test*). Data representative of three independent experiments. **k, l**, Dissemination of *C. rodentium* to the (**k**) spleen and (**l**) liver in *Vip*^*IRES-Cre*^*hM3Dq*^*fl-stop-fl/+*^ mice treated with vehicle or CNO (1mg/Kg, daily) for 4 days post-intragastric infection with 2×10^9^ CFU. Log_10_ Colony Forming Units (CFU) of *C. rodentium* 9 days post-oral innoculation (9 d.p.i.). Dotted line: limit of detection. \*\*\**P*=0.0009 and *\*\*\*\**P<0.0001, (*Mann-Whitney test*). Vehicle: n=11 (Positive for *C. rodentium:* spleen: 6/11, liver: 8/11), CNO: n=9 (Positive for *C. rodentium:* spleen and liver: 9/9). Data representative of two independent experiments. **m**, Survival rates for *C. rodentium*-infected *Vip*^*IRES-Cre*^*hM3Dq*^*fl-stop-fl/+*^ mice treated with vehicle or CNO (1mg/Kg, daily, 1-4 d.p.i.: blue rectangle). Vehicle: n=11, CNO: n=11. Data representative of three independent experiments. \*\*\*\**P*<0.0001 *(Mantel-Cox test*). **n, o**, Bacterial dissemination to the (**n**) spleen and (**o**) liver of *Vip*^*IRES-Cre*^*hM4Di*^*fl-stop-fl/+*^ mice treated with vehicle or CNO (1mg/Kg, daily, 1-4 days post-intragastric infection with 4×10^10^ CFU of *C. rodentium*). Log_10_ CFU at 9 d.p.i. Dotted line: limit of detection. Vehicle: n=8 (Positive for *C. rodentium:* spleen: 8/8, liver: 8/8), CNO: n=7 (Positive for *C. rodentium:* spleen: 1/7, liver: 1/7). \*\*\**P*=0.0006 (spleen) and \*\*\**P*=0.0005 (liver) (*Mann-Whitney test*). Data representative of two independent experiments. **k**,**l**,**n**,**o**: For visualization in the logarithm scale, CFU counts of 0 were attributed a value of 1.

To determine whether direct modulation of VIPergic neurons (inhibition or activation) could affect ILC3 function, we adopted a chemogenetic strategy utilizing mice engineered to express designer receptors exclusively activated by designer drugs (DREADD)^25^ in VIP^+^ cells (Extended Data Figure 6a-c). We expressed inhibitory DREADD (hM4Di) in VIPergic neurons (*Vip*^*IRES-Cre*^*hM4Di*^*fl-stop-fl/+*^ mice) and observed a higher frequency of IL-22-producing CCR6^+^ILC3 in the small and large intestine at 24h following VIPergic inhibition with the DREADD ligand clozapine-N-oxide (CNO, Fig. 2e,f and Extended Data Figure 6d,e). In parallel, there was increased *RegIIIγ* mRNA expression in ileal extracts enriched for IECs after inhibition of VIPergic neurons (Fig. 2g). Conversely, mice expressing the activating DREADD (hM3Dq) in VIPergic neurons (*Vip*^*IRES-Cre*^*hM3Dq*^*fl-stop-fl/+*^ mice) had fewer IL-22^+^ cells among the CCR6^+^ ILC3 in the small and large intestine, and lower *RegIIIγ* mRNA in IEC-enriched ileal extracts following activation of VIPergic neurons (Fig. 2h-j and Extended Data Figure 6f). However, differences were not observed in the frequency of IL-17A or IL-22 production among CD3^+^RORγt^+^ cells (γδT17 and Th17) after activation of VIPergic neurons (Extended Data Fig. 6g-h). Combined, these results indicate that VIPergic neurons modulate intestinal immune homeostasis, acting through VIPR2 on CCR6^+^ ILC3 to inhibit cell activation and IL-22 production.

We next investigated whether alteration of VIPergic neuronal activity contributes to changes of intestinal barrier function during intestinal colonization with an enteropathogen. As previously described^26^, following infection with *Citrobacter rodentium* there was increased *Vip* mRNA expression in the proximal colon and cecum and of VIP amounts in the portal vein, but not in the peripheral blood (Extended Data Fig. 7a-c), suggesting an increase in VIPergic activity in the intestinal tissue. Oral gavage of mice with a relatively low dose of *C. rodentium* (2×10^9^ CFU) is tolerated by mice with an intact immune system. However, DREADD-mediated activation of VIPergic neurons during the first 4 days of infection led to increased bacterial translocation to the spleen and liver (Fig. 2k,l) and resulted in reduced survival (Fig. 2m and Extended Data Fig 7d), despite only a moderate increase in the amount of luminal *C. rodentium* (Extended Data Fig. 7e). Gavage with a high dose of *C. rodentium* (4×10^10^ CFU) resulted in large amounts of bacteria that translocated to the liver and spleen (9d.p.i.) (Fig. 2n,o). DREADD-mediated inhibition of VIPergic neurons provided substantial protection from bacterial breach of the intestinal barrier without affecting the bacterial burden in the feces (Fig 2n,o and Extended Data Fig. 7f). Moreover, treatment with recombinant murine IL-22 reversed the effect of DREADD-mediated activation of VIPergic neurons on mortality and bacterial translocation (Extended Data Fig. 7g,h). In addition, *RORc(t)*^*Cre*^*Vipr2*^*fl/fl*^ mice exhibited enhanced protection from bacterial breach of the intestinal barrier after gavage with a high dose of *C. rodentium* (4×10^10^ CFU) when compared with *RORc(t)*^*Cre*^*Vipr2*^*+/+l*^ mice (Extended Data Figure 7i,j). Activation of VIPergic neurons during the events that follow infection with enteropathogens may therefore be an important contributor to intestinal barrier breakdown, which could be mitigated by inhibition of VIPergic signaling in ILC3.

These results suggested the existence of physiological signals that promote activation of VIPergic neurons during homeostasis, with consequent tonic inhibition of CCR6^+^ ILC3. Food ingestion has been reported to rapidly signal VIP release in the intestine^4^. Indeed, we observed a greater amount of VIP in the ileum of mice sampled during the vivarium’s dark-phase as compared to the light-phase, which correspond to periods of food consumption and resting, respectively (Extended Data Fig. 8a). Due to the *ad libitum* feeding scheme, approximately 15% of daily food intake occurs during the light-phase/resting period^27^. To dissect the effect of food intake on the VIPergic immune inhibitory axis and to reduce the noise created by feeding during the resting period, we restricted food availability for two weeks to alternating 12h cycles of feeding and fasting. To dissociate the effects of light/dark cycles from time of feeding, food was available to a group of mice only during the dark-phase (night-fed mice, ZT12-ZT0/6PM-6AM), while another group had food available only during the light-phase (day-fed mice, ZT0-ZT12/6AM-6PM). Using this scheme, we observed more VIP in the portal vein, but not in the peripheral blood, after 6h of food availability when compared to mice fasted for 6h, regardless of whether mice were fed during the light phase (ZT 6) or during the dark phase (ZT 18) (Extended Data Fig. 8b,c). In turn, the frequency of IL-22^+^ CCR6^+^ ILC3 was reduced during the periods of food consumption (at 6h of feeding) relative to fasting periods (fasted for 6h), independently of light-dark cycles (Fig. 3a,b).

**Figure 3.**
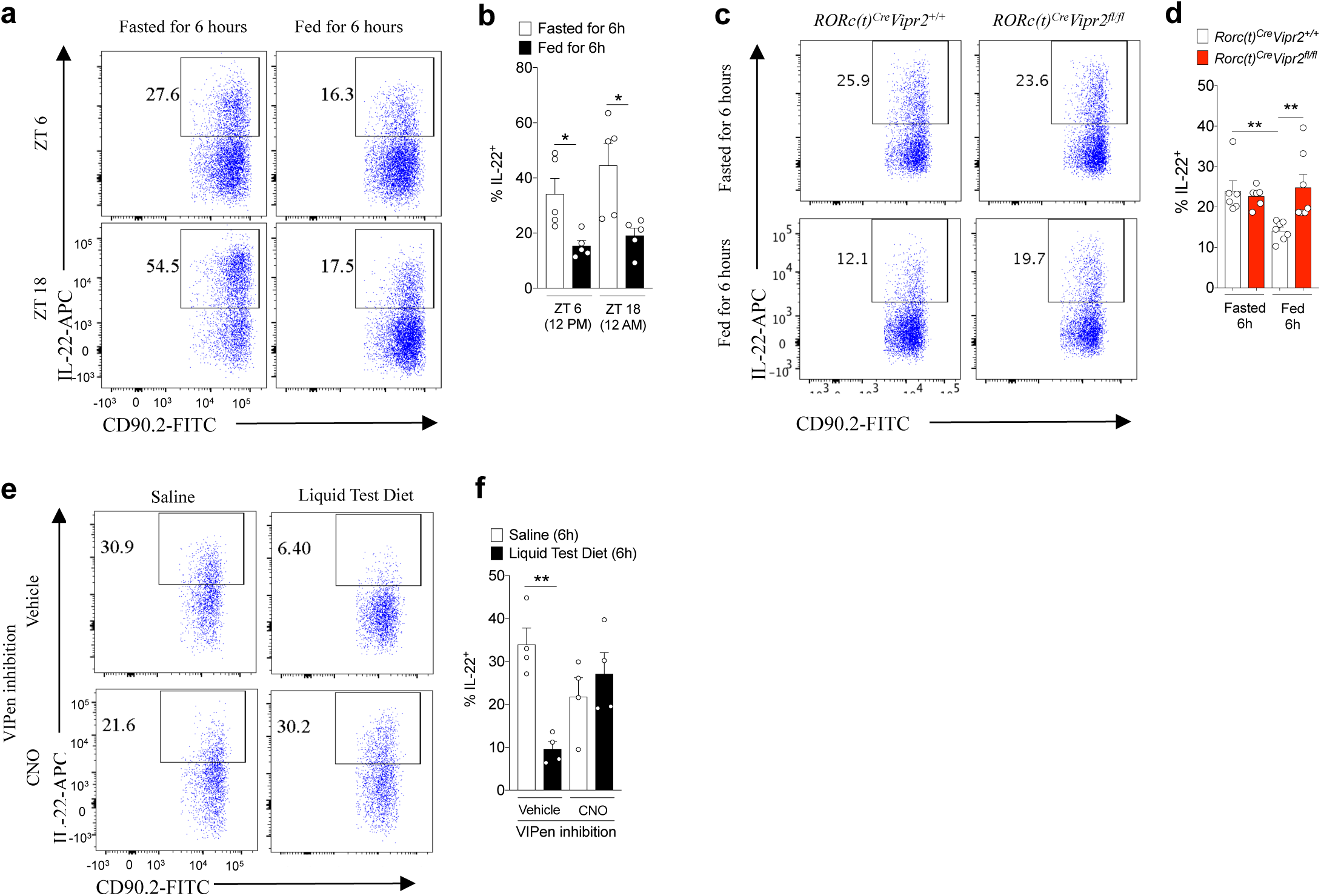
Feeding reduces IL-22 production by CCR6^+^ILC3 through activation of VIPergic neurons. **a, b**, Representative FACS plot (**a**) and summaries (**b**) indicating IL-22 expression among CCR6^+^ ILC3 from the ileum of mice at 6h after feeding or fasting, at ZT6 and ZT18. N=5, \**P*<0.05 (*t-test*). Data are representative of two independent experiments. **c, d**, Representative FACS plot (**c**) and summaries (**d**) indicating IL-22 expression by CCR6^+^ ILC3 from *RORc(t)*^*Cre*^*Vipr2*^+/+^ or *RORc(t)*^*Cre*^*Vipr2*^fl/fl^ mice at 6h after fasting or 6h after feeding (ZT 18). \*\**P*<0.01 (*paired t-test*). Data shown are pooled from two independent experiments and representative of three independent experiments. **e, f**, Representative FACS plot (**e**) and summaries (**f**) indicating IL-22 expression in CCR6^+^ ILC3 from the ileum of *Vip*^*IRES-Cre*^*hM4Di*^*fl-stop-fl/+*^ mice (DREADD for VIPergic inhibition). Mice were treated with vehicle or CNO (1mg/Kg) and 30 minutes later were fed by intragastric administration of saline (0.4 mL each 45 min, for 6 h) or Liquid Test Diet (500 mg/mL, 0.4mL each 45 minutes, for 6 h). N=4, \*\**P*<0.01 (*t-test*). Data shown are representative of three independent experiments.

Using the same feeding scheme, we observed among CCR6^+^ ILC3 from *RORc(t)*^*Cre*^*Vipr2*^*+/+*^ mice a reduced frequency of IL-22^+^ cells at 6h after start of food consumption (ZT 18) when compared to mice fasted for 6h (ZT 6) (Fig 3c,d). In contrast, there was no difference in frequency of IL-22^+^ cells in the CCR6^+^ ILC3 from *RORc(t)*^*Cre*^*Vipr2*^*fl/fl*^ littermates with or without feeding. The same results were observed with mixed bone marrow chimeric mice (Extended Data Figure 8d,e). This is consistent with VIPR2 signaling-dependent inhibition of IL-22 production in CCR6^+^ ILC3 following food intake. This conclusion was further supported by the observed reduction in the frequency of IL-22^+^ CCR6^+^ ILC3 at 6h following intragastric delivery of a liquid test diet, but not control saline, during the light-phase period (ZT1-ZT7) (Fig 3e,f). Furthermore, DREADD-mediated inhibition of VIPergic neurons abrogated the effect of the test diet on ILC3 (Fig. 3e and 3f). These results suggest a fast and dynamic temporal control of intestinal CCR6^+^ ILC3 function promoted by food intake-mediated activation of VIPergic signaling and VIPR2.

Because IL-22 acts on intestinal epithelial cells to regulate barrier functions, including their interactions with commensal microbiota, we next examined whether the neuroimmune inhibitory circuit promoted by food consumption influences host-microbial interactions. There was an IL-22-dependent elevation of *RegIIIγ* mRNA expression in IEC-enriched ileal extracts from mice fasted for 12h as compared to those fed during that time frame (Fig. 4a). In parallel, food consumption had a striking effect on the morphology of the ileal epithelium-associated segmented filamentous bacteria (SFB) (Fig. 4b-c and Extended Data Fig. 9a-c). SFB attached to IECs had long segmented filaments after 12h of food consumption, independently of light/dark cycles. However, in the ileum of mice fasted for 12h, independently of light-cycle, epithelium-associated SFB had few segments and were stubble-like. A strong effect of feeding/fasting cycles was also observed in the composition of the fecal microbiota, independently of light-cycle (Extended Data Fig. 9d,e). Indeed, there was an increase in the *Firmicutes/Bacteroidetes* ratio in the feces of mice after 12h of feeding compared to 12h of fasting.

**Figure 4.**
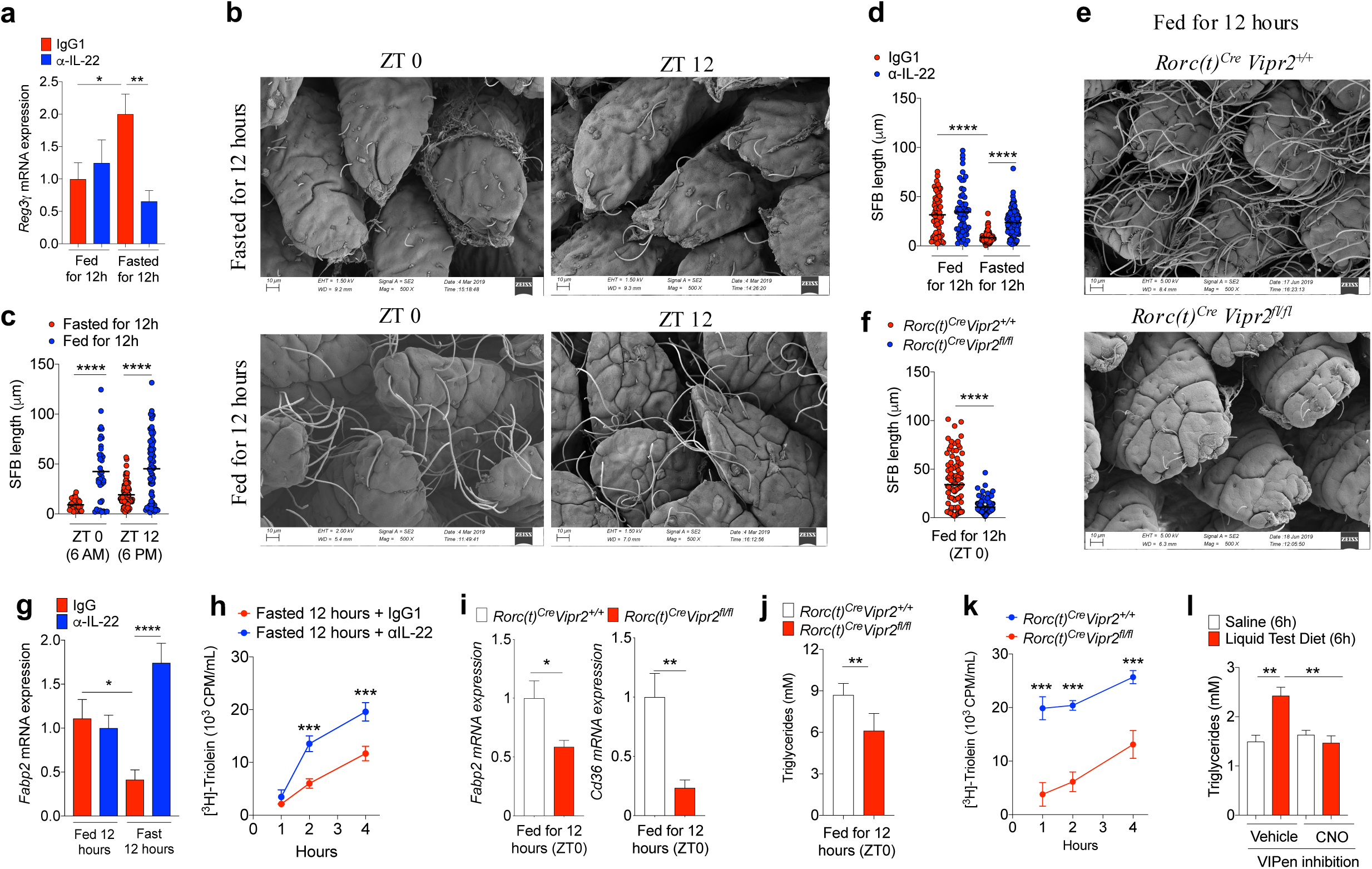
Dynamic regulation of commensal bacterial growth and lipid absorption by feeding-dependent modulation of the VIP-VIPR2-IL-22 axis. **a**, Normalized *RegIIIγ* mRNA expression in IECs from mice treated with IgG or *α*-IL-22 (10mg/Kg) during the feeding period (Fed for 12h: Treated from ZT 12->ZT 0) or during the fasting period (Fasted for 12h: Treated from ZT 0->ZT 12). Data pooled from 2 independent experiments: Fed IgG1: N=8, Fed *α*-IL-22: N=8, Fasted IgG1: N=8, Fasted *α*-IL-22: N=7, \**P*<0.05 and \*\**P*<0.01 (*t-test*). **b, c**, Representative scanning electron microscopy (SEM) images (**b**) of epithelium-associated commensal SFB in the ileum of mice 12h after fasting or feeding at the end of the dark-phase (ZT 0) and the light-phase (ZT 12) and (**c**) measurements of SFB filament lengths. N=3, data representative of 2 independent experiments,\*\*\*\**P*<0.001 (*t-test*). **d**, SFB length in the ileum of mice treated with IgG or *α*-IL-22 (10mg/Kg) during the feeding period (Fed for 12h: ZT 12->ZT 0) or during the fasting period (Fast for 12h: ZT 0->ZT 12). N= 3, data representative of 2 independent experiments,\*\*\*\**P*<0.001 (*t-test*). **e**,**f**, Representative **(e)** SEM images of epithelium-associated SFB in the ileum of *RORc(t)*^*Cre*^ *Vipr2*^*+/+*^ and *RORc(t)*^*Cre*^ *Vipr2*^*fl/fl*^ mice fed for 12h and **(f)** measurement of bacterial filament lengths. N= 3, \*\*\*\**P*<0.001 (*t-test*). **g**, Normalized *Fabp2* mRNA expression in IEC-enriched ileal extracts from mice treated with IgG or α-IL-22 (10mg/Kg) during the fasting period (Fasted for 12h: Treated from ZT 0->ZT 12). Data representative of 3 independent experiments: Fed IgG1: N=8, Fed *α*-IL-22: N=8, Fast IgG1: N=8, Fed *α*-IL-22: N=7. \**P*<0.05 and \*\*\*\**P*<0.0001 (t-student). **h**, Plasma ^3^H CPM (counts per minute) in 12h fasted mice after gavage with ^3^H-triolein (2uCi/mice in 200ul of 20% Intralipid). Mice were treated with IgG or *α*-IL-22 (10mg/Kg) during the fasting period (Fasted for 12h: Treated from ZT 0->ZT 12). Data representative of 2 independent experiments. N=4, \*\*\**P*<0.001 (*two-way ANOVA*). **i, j**, Normalized *Fabp2* and *Cd36* mRNA expression in IEC-enriched ileal extracts from (**i**) and plasma triglyceride content (**j**) in 12h fed *RORc(t)*^*Cre*^ *Vipr2*^*+/+*^ (N=5) and *RORc(t)*^*Cre*^ *Vipr2*^*fl/fl*^ (N=4) mice. *\*P*<0.05 and \**\*P*<0.01 (*t-test*). **k**, Plasma ^3^H CPM in 12h fed *RORc(t)*^*Cre*^ *Vipr2*^*+/+*^ (N=3) and *RORc(t)*^*Cre*^ *Vipr2*^*fl/fl*^ (N=3) mice after gavage with ^3^H-triolein. Data representative of 2 independent experiments,\*\*\**P*<0.001, (*two-way ANOVA*). **l**, Plasma triglyceride content in *Vip*^*IRES-Cre*^*hM4Di*^*fl-stop-fl/+*^ mice (DREADD for VIPergic inhibition) after gavage with Saline (0.4 mL/each 45 minutes, for 6 hours) or Liquid Test Diet (500 mg/mL, 0.4mL/each 45 minutes, for 6 hours). Vehicle: N=5, CNO: N=4. Data representative of 2 independent experiments, \*\**P*<0.01, (*t-test*).

Short-term blockade with *α*-IL-22 not only reduced *RegIIIγ* mRNA expression in the ileal extracts enriched for IECs of mice during fasting, but also prevented control of SFB growth upon food restriction (Fig. 4d). In *RORc(t)*^*Cre*^*Vipr2*^*fl/fl*^ mice, which have increased IL-22 production, SFB failed to form segmented filaments even after 12h of food consumption (Fig. 4e,f and Extended Data Figure 10a). Moreover, we observed a reduced *Firmicutes/Bacteroidetes* ratio in the ileal fecal matter of *RORc(t)*^*Cre*^*Vipr2*^*fl/fl*^ mice after 12h feeding compared to *RORc(t)*^*Cre*^*Vipr2*^*+/+*^ mice (Extended Data Figure 10b-e). These results suggest that VIPergic signaling regulate commensal microbiota by modulating cryptopatch-associated ILC3 in response to food consumption.

As IL-22 was recently shown to regulate lipid metabolism^8^, at least in part by controlling genes involved in lipid transport, we examined the role of the VIPergic-ILC3-IL-22 circuit in lipid absorption. Mice were adapted for 2 weeks to 12h restricted feeding during light or dark cycles and were then gavaged with ^3^H-triolein following feeding or fasting. Mice that had been fed for 12h absorbed more of the triglyceride than mice fasted for 12h, although there was some effect of circadian regulation (light/dark adaptation) (Extended Data Fig. 11a,b). Expression of mRNAs for proteins associated with fatty-acid/lipid uptake and transport (e.g. *Fabp2*) was lower in IEC-enriched ileal extracts from mice that were food-restricted for 12h compared to fed mice (Fig. 4g), in inverse relationship to *RegIIIγ* mRNA (Fig 4a). Moreover, short-term treatment with *α*-IL-22 during the fasting period increased ^3^H-triolein absorption and the expression of lipid transporters in ileal extracts enriched for IECs (Fig 4g,h).

The effect of food intake on IEC-dependent lipid absorption was dependent on VIPergic signaling of ILC3, since expression of *Fabp2* and *Cd36* mRNAs, ^3^H-triolein absorption, and concentrations of circulating plasma triglycerides were substantially reduced in *RORc(t)*^*Cre*^*Vipr2*^*fl/fl*^ mice when compared to *RORc(t)*^*Cre*^*Vipr2*^*+/+*^ littermates after 12h of feeding (ZT 0) (Figures 4i-k). Moreover, *RORc(t)*^*Cre*^*Vipr2*^*fl/fl*^ mice had reduced weight compared to their *RORc(t)*^*Cre*^*Vipr2*^*+/+*^ littermates (Extended Data 11c,d), similarly to transgenic mice that constitutively express high levels of IL-22^28^. Finally, continuous intragastric delivery of a liquid test diet for 6h during the light phase period led to an increase in serum triglycerides, which was blocked by DREADD-mediated inhibition of VIPergic neurons (Fig. 4l).

These results reveal an important neuro-immune circuit in which feeding-activated VIPergic neurons antagonize microbiota-dependent CCR6^+^ ILC3 function, resulting in reduced IL-22 production and increased efficiency of lipid absorption (Extended Data Figure 11e,f). The benefit of greater nutrient acquisition comes at the expense of reduced IL-22-promoted antimicrobial functions, illustrated by susceptibility to enteropathic bacteria, rapid growth of epithelial-associated commensal SFB, and broader changes in the luminal commensal composition (increased *Firmicutes/Bacteroidetes* ratio). Whether the host derives benefit from the reduced anti-microbial activity is unclear, although it may enable bacteria-dependent generation of essential metabolites^29^ and vitamins that would be readily absorbed. It will be of interest to determine in the future the global metabolic consequences following chronic activation of ILC3/IL-22 in the absence of VIPR2 signaling. Since enteric glial cells (EGCs) are reported to secrete factors that act as activators of ILC3 functions^10^, it will additionally be important to determine whether VIPergic neurons and EGCs coordinate antagonistic signals to promote efficient modulation of ILC3-mediated intestinal barrier functions, and also how neural development-associated factors expressed by ILC3 (e.g. *Cntn1, Sema3d, Bmp2*) contribute to formation of neuro-immune cell units in cryptopatches/ILFs.

The mechanism by which the presence of food in the intestinal lumen is sensed and promotes VIPergic activation is still unclear, and microbial^30,31^, nutritional^4^ or mechanical stimuli^32^, acting peripherally or by way of the central nervous system, may be responsible for this. The VIP system has been implicated in circadian regulation centrally, as VIPR2 is required for expression of the core clock genes in the suprachiasmatic nuclei of the hypothalamus, and in the periphery, with circadian oscillation of IL-5 and IL-13 production by ILC2 and feeding-associated eosinophil recruitment in lung and gut mucosa^33,34^. While our study highlights the importance of food intake in regulating VIPergic neurons, there is also a light-dependent contribution to the efficiency of lipid absorption, as there was only a partial effect when the feeding cycle was reversed. We therefore cannot rule out an additional role for central or peripheral clock functions^35^ independent of the enteric VIP-ILC3 circuit.

Our results suggest that disturbances in feeding schedules could reduce intestinal barrier functions and promote dysbiosis and at the same time contribute to metabolic inbalance by chronic activation of VIPergic neurons. Moreover, enteropathogens may hijack this neuro-immune circuit and reduce intestinal barrier functions, facilitating intestinal colonization. In these scenarios, VIPR2 inhibitors may be valuable therapeutic tools to reduce lipid absorption or enforce the barrier during acute gastrointestinal infections.

## Acknowledgements

We thank members of the Littman lab and Juan J. Lafaille for valuable discussion. We thank S.Y. Kim and the NYU Rodent Genetic Engineering Laboratory (RGEL) for generating the mutant mice, Cindy Loomis and the Experimental Pathology Research Laboratory of NYULMC for histology of small intestine samples, and Adriana Heguy and the Genome Technology Center (GTC) for RNA sequencing. We also thank F-X. Liang, J. Sall, C. Petzold and K. Dancel at the Microscopy Laboratory Core for timely preparation of the scanning electron microscopy images, and M. Cammer and Y. Deng for the help with optical microscopy. The Microscopy Core and the GTC are partially supported by NYU Cancer Center Support Grant NIH/NCI P30CA016087 at the Laura and Isaac Perlmutter Cancer Center, S10 RR023704-01A1 and NIH S10 ODO019974-01A1. The Experimental Pathology Research Laboratory is supported by National Institutes of Health Shared Instrumentation grants S10OD010584-01A1 and S10OD018338-01. This study benefitted from data assembled by the ImmGen Consortium. This work was supported by the Pew Latin American Fellows program (J.T.), the Helen and Martin Kimmel Center for Biology and Medicine (D.R.L.); and National Institutes of Health grant R01DK103358 (D.R.L.). D.R.L. is an Investigator of the Howard Hughes Medical Institute.

## Author Contributions

J.T. and D.R.L. designed the study and analyzed the data. J.T. performed the experiments with assistance from P.H. and D.L., H.N. contributed to the feeding experiments, and L.K. performed the bioinformatics analyses. J.T. and D.R.L. wrote the manuscript. D.R.L. supervised the research.

## METHODS

### Mice

C57Bl/6 mice were obtained from Taconic Farm. All transgenic mice were bred and maintained in the animal facility of the Skirball Institute (New York University School of Medicine) in specific pathogen-free conditions. *hM3Dq*^*fl-stop-fl*^ mice (CAG-LSL-Gq-DREADD, Jax #026220), *hM4Di*^*fl-stop-fl*^ mice (CAG-LSL-Gi-DREADD, Jax #026219), *VIP*^*IRES-Cre*^ mice (B6J.Vip-IRES-Cre, Jax #031628), *tdTomato*^*f/-stop-fl*^ (CAG-tdTomato, JAX #007914) and CD45.1 mice (*B6.SJL-Ptprca Pepcb/BoyJ*, Jax# 002014) were purchased from Jackson Laboratories. *RORc(t)*^EGFP/+^ and *RORc(t)*^Cre^ mice were generated in our laboratory and previously described^14,36^. BAC-transgenic *Rorc(t)-Gfp*^*TG*^ were generated by G. Eberl’s lab^37^ and used for evaluation of *Vipr2* mRNA expression in different populations of RORγt^+^ lymphoid cells purified by FACS sorting. *Vipr2*^*-/-*^ mice (Jax# 031332) were purchased from Jackson Laboratories and were maintained in a CD45.2 background or were bred with CD45.1 (*B6.SJL-Ptprca Pepcb/BoyJ*) mice, which subsequently generated *Vipr2*^*-/-*^ CD45.1/2 mice and WT CD45.1 littermates. Conditional VIPR2 knockout (*Vipr2*^*fl/fl*^) mice were generated using CRISPR-Cas9 technology as described below. Mice in all experiments were 6–12 weeks old at the starting point of treatments and all were colonized with SFB. All animal procedures were performed in accordance with protocols approved by the Institutional Animal Care and Usage Committee of New York University School of Medicine or the NIAID as applicable.

### Generation of RORc(t)^Cre^ Vipr2^fl/fl^ mice

Mice carrying *loxP* sites flanking exons 3 and 4 of the Vipr2 gene (*Vipr2*^*fl/fl*^) were generated using published CRISPR/Cas9 protocols^38,39^ at the NYU School of Medicine’s Rodent Genetic Engineering Laboratory. Briefly, guide RNAs targeting regions upstream of exon 3 (gRNA_16 sequence: GAAATCTCACAACAGATTCG) and downstream of exon 4 (gRNA_23 sequence: TCTCCTCAGAAGCATCGAAT) were designed using the Crispr guide design software (http://crispr.mit.edu/). gRNA recognition sequences were cloned into the pX330 vector (a gift from Dr. Feng Zhang, Addgene #42230), using oligos with a T7 promoter containing the gRNA template that were chemically synthetized by IDT (Integrated DNA Technologies). The products of PCR-amplified T7-gRNA were used as templates for in vitro transcription (MEGAshortscript T7 kit, Thermo Fisher Scientific). The gRNAs were purified using the MEGAClear Transcription Clean-up kit (Thermo Fisher Scientific). Two donor templates (ssDNA_1 and ssDNA_2) with 200bp each were chemically synthetized by IDT. The donor templates contained 60bp homology arms flanking a cassete containing a loxP sequence and XhoI or SalI restriction sites, located in the original PAM sequence (mutated PAM). Namely, ssDNA_1 (XhoI) for the region upstream exon 3: (GGATGGCATTTACATAGGACCCCATCCCAGTGGCTGCTCAGAAGAGCACTCACTCCTTATCCctcgagataacttcgtataatgtatgctatacgaagttatCCGAATCTGTTGTGAGATTTCGAGAACTCATAAGGACTGATAAGGCCACACAACTTGAGC), and ssDNA_2 for the region downstream exon 4 (TGATTTCTCCTAGGTCACACTCAGGGAGCATTTCCAGACACTGGAAAACTCCTGAGGCCCgtcgacataacttcgtataatgtatgctatacgaagttatATTCGATGCTTCTGAGGAGACTATAATTAAACCCTGCCTGTGTGAGGCATGGCTTCTGAT). Mice were generated by injection of a mixture of mammalian optimized Cas9 mRNA (100 ng/μl, TriLink Biotechnologies), purified gRNA_16 and gRNA_23 (50 ng/μl, each) and donor templates ssDNA_1 and ssDNA_2 (50ng/ul, each) in injection buffer (10 mM Tris, pH 7.5; 0.1 mM EDTA) into the cytoplasm of C57BL/6J embryos in accordance with standard procedures approved by the IACUC at the NYU School of Medicine. Female CD-1 mice (Charles River) were used as foster mothers. F0 mice were genotyped and sequenced (Sanger sequencing) to identify mice homozygous for both loxP insertions. Founders bearing loxP insertions were then backcrossed at least one time to wild-type C57BL/6J mice generating the *Vipr2*^*fl/fl*^ mice. For the generation of *RORc(t)*^*Cre*^ *Vipr2*^*fl/fl*^ mice, *Vipr2*^*fl/fl*^ mice were crossed with *RORc(t)*^Cre^ mice for the generation of *RORc(t)*^Cre^ *Vipr2*^*fl/fl*^ and *RORc(t)*^Cre^ *Vipr2*^*+/+*^ littermates.

### VIPergic neuronal activation/inhibition using DREADDs

To perform chemogenetic activation or inhibition of VIPergic neurons, we bred *VIP*^*IRES-Cre*^ homozygous mice to *hM3Dq*^*fl-stop-fl*^ mice (DREADD for activation) or *hM4Di*^*fl-stop-fl*^ mice (DREADD for inhibition), generating *VIP*^*IRES-Cre*^*hM3Dq*^*fl-stop-fl*^ (VIPergic activation) and *VIP*^*IRES-Cre*^ *hM3Dq*^*fl-stop-fl*^*hM4Di*^*fl-stop-fl*^ mice (VIPergic inhibition). To perform 24h activation of the DREADDs, mice were treated with Clozapine-N-Oxide (CNO, 1mg/Kg intraperitoneally, TOCRIS) each 12h. Our experiments with *VIP*^*IRES-Cre*^ only mice revealed that these dose of CNO treatment does not affect ILC3 function. To perform activation of the DREADDs during *Citrobacter rodentium* infection, mice were treated daily with CNO (1mg/Kg, intraperitoneally) from day 1 to day 4 after infection.

### Food restriction protocol

For a period of two weeks, food was made available to mice only for 12h per daily cycle. Mice were kept in two different regimens: Dark-phase fed mice, with food being available between 6 PM – 6AM (ZT 12->ZT 0); and light-phase fed, with food being available between 6 AM – 6PM (ZT 0->ZT 12). To avoid littering, at the beginning of the fasting period of each regimen, mice were transferred to a clean cage containing alpha cellulose clean bedding (Shepherd’s™ALPHA-dri). Mice were provided with free access to water.

### Gavage of Liquid Test Diet

Dry powder micro stabilized rodent liquid diet (Test Diet, LD 101) was blended (mechanical blender) vigorously for 30 seconds in saline (NaCl 0.9%) at 0.5g/mL. Mice were gavaged with 400ul of this solution using a polyurethane feeding tube (16ga x 38mm, FTPU-16-38-50, INSTECH) every 45 min for 6 h.

### Generation of bone marrow VIPR2^+/+^/VIPR2^-/-^ chimeric reconstituted mice

Bone marrow mononuclear cells were isolated from CD45.1 *VIPR2*^*+/+*^ and CD45.2 (or CD45.1/2) *VIPR2*^*-/-*^ mice by flushing the long bones. Red blood cells were lysed with ACK Lysing Buffer and the remaining cells were resuspended in PBS for injection in at a 1:1 ratio (WT:VIPR2 KO). 4×10^6^ cells were injected intravenously into 6 week old CD45.1/2 (CD45.2) mice that were irradiated 4h before reconstitution using 1000 rads/mouse (2×500rads, at an interval of 3h, at X-RAD 320 X-Ray Irradiator). To deplete intestinal ILC3, one day after the bone marrow transfer, mice were treated with InVivoMAb anti-mouse Thy1.2 (200ug/mice for 4 consecutive days, Clone 30H12, BioXCell). Experiments were performed 6-7 weeks after the last treatment with α-Thy1.2.

### Radioactively labeled triglyceride absorption assay

Plasma ^3^H-CPM (counts per minute) was measured 1-4h after gavage with ^3^H-Triolein– containing lipid^40^. Briefly, mice were injected with poloxamer 407 (1g/Kg, i.p., Sigma, #16758). After 30 minutes, mice were gavaged with a mixture of 2μCi ^3^H-Triolein in 200ul of 20% intralipid oil emulsion. Blood samples were collected and diluted in Liquid Scintillation Counting cocktail (Ultima Gold) and measured using a Scintillation counter (Beta Counter MicroBeta^2^ System, Perkin Elmer).

### IL-22 in vivo Blockade

For experiments with IL-22 blockade, mice were injected with monoclonal α-IL-22 (Clone 8E11, 250μg/mouse, generously provided by Tangsheng Yi, Genentech), 12 hours before sample collection. Control groups received mIgG1 (inVivoMAb, BioXCell).

### C. rodentium mediated colon inflammation

*C. rodentium* strain DBS100 (ATCC51459; American Type Culture Collection) was grown at 37°C in LB broth to OD600 reading between 0.5 and 0.7. *VIP*^*IRES-Cre*^*hM3Dq*^*fl-stop-fl*^ (VIPergic activation) and C57BL/6 mice were inoculated with 200 μl of a bacterial suspension (2×10^9^ CFU) by oral gavage. *VIP*^*IRES-Cre*^ *hM3Dq*^*fl-stop-fl*^*hM4Di*^*fl-stop-fl*^ mice (VIPergic inhibition) and *RORc(t)*^*Cre*^ *Vipr2*^*fl/fl*^ were inoculated with 200 μl of a bacterial suspension (3-4×10^10^ CFU) by oral gavage. For DREADD experiments, CNO treatment started at 1 day post-infection (d.p.i.) until 4 d.p.i. Mice were followed the next 12 days post-infection (d.p.i.) to measure survival rate. At 9 d.p.i. fecal pellets were collected and the mice dissected to harvest spleen and liver. Samples were weighted and minced on sterile deionized water with Triton 0.1% and filtered on a 70μm cell strainer. The filtered samples were used to measure *C. rodentium* burden with serial dilutions (triplicates) on MacConkey agar plates.

### Immunofluorescence and Confocal Microscopy

Slide section staining: Small intestines from perfused SFB^+^ *Rorc*(t)^EGFP/+^ or *VIP*^*IRES-Cre*^*tdTomato*^*fl-stop-fl*^ mice were Swiss-rolled, fixed for 4h in 4% paraformaldehyde, incubated overnight in 30% sucrose at 4°C, and frozen in embedding medium for frozen specimens (O.C.T, Tissue-Tek, Sakura). Tissue was cut into 30-70 µM sections, blocked in PBST 0.5% (0.5% Triton X-100, 10% normal donkey serum) for 1 h, and incubated overnight at 4°C using the following antibodies: α-Vasoactive Intestinal Peptide (1:1000, rabbit polyclonal, 20077, Immunostar), α-Tyrosine Hydroxilase (1:50, rabbit polyclonal, AB152, Millipore), α-Substance P (1:3000, rabbit polyclonal, 20064 Immunostar), α-GFP Alexa Fluor 488 (1:500, clone: FM264G, Biolegend), α-TCRβ Brilliant Violet 421 (1:50, Biolegend), α-β-3-Tubulin Alexa Fluor 594 or Alexa Fluor 488 (1:500, clone:AA10, Biolegend). Tissue was washed and when needed incubated with secondary fluorescently labeled antibodies (Donkey Anti-Rabbit Pacific Blue or Alexa Fluor 647) for 2h before nuclear staining with Draq-7 (R&D Systems) or 4′,6-diamidino-2-phenylindole (DAPI, ThermoFisher).

Whole mount stainings: Ileum was collected from perfused *VIP*^*IRES-Cre*^*hM3Dq*^*fl-stop-fl*^ mice, flat opened and cleared of fecal material. The flat tissue was fixed overnight in 4% paraformaldehyde at 4°C. Whole tissue was cut in sections of 4cm, blocked in PBST 0.5% (0.5% Triton X-100, 10% normal donkey serum) overnigh at 4°C, and incubated for 48h at 4°C using the following antibodies: α-HA Alea Fluor 647, for staining of HA-tagged hM3Dq, (16B12, 1/300, Biolegend) and α-cFOS (1/100, 9F6, Rabbit, Cell, Signaling #2250). Tissue was washed for 24h at 4°C and incubated with secondary fluorescently labeled antibodies (Donkey Anti-Rabbit Alexa Fluor 488, Thermo Fisher) for 24h at 4°C. Tissue was washed for 24h before nuclear staining with 4′,6-diamidino-2-phenylindole (DAPI, ThermoFisher). Images were acquired using a Zeiss LSM 710 confocal (Carl Zeiss). The imaging data were processed and analyzed using Image J software (NIH, Bethesda, MD). Imaris software version 9.0.1 (Bitplane; Oxford Instruments) was used to generated reconstructed 3D images.

### Isolation of Lamina Propria Lymphocytes (LPLs) from the small intestine

Whole small intestine, proximal small intestine (14cm), ileum (distal 14cm of the small intestine) or the large intestine were dissected from mice. Unless indicated, all the dissections were performed between ZT2-ZT3. Mesenteric fat tissue and Peyer’s patches were carefully removed from these tissues. Intestinal tissue was opened and extensively cleaned of fecal matter. Following, this tissue was sequentially treated with HBSS 1X (1 mM DTT) at 37°C for 10 min with gentle shaking (200rpm), and twice with 5 mM EDTA at 37°C for 10 min to remove epithelial cells. The EDTA fraction (ileal extracts enriched for IECs) was filtered using a 100uM strainer, centrifuged and suspended in Trizol for further RNA isolation. The remaining tissue was then minced with a scissor and dissociated in RPMI containing 10% FBS, Dispase (0.05 U/ml; Worthington), collagenase (1 mg/ml collagenase II; Roche) and DNase I (100 μg/ml; Sigma) at constant shaking at 37°C for 45 min (175rpm). The digested tissue was then filtered through a 70μm strainer to remove large debris. Viable Lamina Propria Lymphocytes (LPLs) were collected at the interface of a 40%/80% Percoll/RPMI gradient (GE Healthcare).

### Isolation of ILC3, γδT17 and Th17 cells from the small intestine

CCR6^+^ and CCR6^neg^ ILC3

(DAPI^neg^CD3^neg^CD11b^neg^CD11c^neg^CD14^neg^CD19^neg^TCRβ^neg^TCRγ^neg^NK1.1^neg^KLRG1^neg^CD127^+^ CD90.2^+^), γδT17

(DAPI^neg^CD11b^neg^CD11c^neg^CD8^neg^D14^neg^CD19^neg^NK1.1^neg^TCRγ^+^TCRβ^neg^GFP^+^CD3^+^) and Th17 (DAPI^neg^CD11c^neg^CD8^neg^CD14^neg^CD19^neg^NK1.1^neg^TCRγ^neg^TCRβ^+^GFP^+^CD3^+^) were isolated from small intestine of the BAC-transgenic *Rorc(t)-Gfp*^*TG*^ mice using the ARIA II FACS Sorter (sorted through a 70μM nozzle, BD Biosciences) and resuspended in TRIZOL for further RNA isolation. CCR6^+^ ILC3 were also isolated from *VIPR2*^*+/+*^*/VIPR2*^*-/-*^ chimeric reconstituted mice for further RNA isolation.

### ILC3 in vitro cell culture

CCR6^+^ and CCR6^neg^ ILC3

(DAPI^neg^CD3^neg^CD11c^neg^CD14^neg^CD19^neg^TCRβ^neg^TCRγ^neg^NK1.1^neg^KLRG1^neg^CD127^+^CD90.2^+^) were isolated from small intestine LPLs of C57BL/6 mice using the ARIA II FACS Sorter (BD Biosciences). ILC3 were cultured at 37°C in flat bottom 96 well plates (10^4^ cells/well) in RPMI supplemented with 10% heat-inactivated FBS (Hyclone), 50 U penicillin-streptomycin (Hyclone), 2 mM glutamine (Hyclone), 10mM HEPES (Hyclone), 1mM sodium pyruvate (Hyclone) and 50 μM β-mercaptoethanol (Gibco). ILC3 were stimulated with IL-23 (100-300 pg/mL, R&D systems) and/or VIPR2 ligands (BAY-559837: 1-100nM, and VIP: 1nM-1uM, TOCRIS) for 16h (37°C), washed, incubated in for 4h at 37°C in RPMI with 10% FBS, phorbol 12-myristate 13-acetate (PMA) (50 ng/ml; Sigma), ionomycin (500 ng/ml;Sigma) and Golgi Plug (BD Bioscience), and stained for membrane extracellular markers in Staining Buffer (PBS FBS 2% EDTA 5mM) and for intracellular markers using Cytofix/Cytoperm buffer set following manufacturer’s protocol (BD Biosciences). Acquisition of cytometric parameters was performed on an LSRII (BD Biosciences). All data were analyzed using FlowJo Software Version 10 (Tree Star).

### Antibodies for intracellular staining and flow cytometry

Single cell suspensions were pelleted and resuspended with surface-staining antibodies in PBS FBS 2% EDTA 1mM. Staining was performed for 20-30min on ice. Live/dead fixable blue (ThermoFisher) was used to exclude dead cells. The following monoclonal antibodies were purchased from eBiosciences, BD Pharmingen or BioLegend: CD3, CD45.1, CD45.2, TCRβ, CD11c, CD14, CD19, KLRG1, TCRγ, RORγ/γt, NK1.1, CD127, CD90.2, CCR6, Sca-1, IL17A and IL-22. Analysis of ex-vivo cytokine production in ILC3 was performed in unstimulated cells (unless indicated), with LPLs incubated for 4h at 37C in RPMI with 10% FBS and GolgiPlug (BD). Cells were then stained for surface markers and anti-CD16/anti-CD32 before fixation and permeabilization, and then subjected to intracellular cytokine staining for IL-22 and IL-17A according to the manufacturer’s protocol (Cytofix/Cytoperm buffer set from BD Biosciences). For analysis of ex-vivo cytokine production by Th17/γδT17 cells were incubated for 4 hours at 37C in RPMI with 10% FBS, phorbol 12-myristate 13-acetate (PMA) (50 ng/ml; Sigma), ionomycin (500 ng/ml;Sigma) and GolgiStop (BD). After surface and live/dead staining, cells were treated using the the FoxP3 staining buffer set from eBioscience according to the manufacturer’s protocol. Intracellular stains were prepared in 1X eBioscience permwash buffer containing anti-CD16/anti-CD32, normal mouse IgG (conc), and normal rat IgG (conc). Staining was performed for 30-60min on ice. Flow cytometric analysis was performed on an LSR II (BD Biosciences) or an Aria II (BD Biosciences) and analyzed using FlowJo software version 10 (Tree Star).

### Blood collection

Peripheral and portal vein blood were collected under general anesthesia (Ketamine 100mg/Kg, Xylazine 15mg/Kg). Peripheral blood was collected through orbital venous plexus bleeding with a glass capillary in a tube containing EDTA (25mM) as an anticoagulant. Plasma was collected after centrifugation of the collected sample and frozen until processing. Surgery was performed to collect blood from the portal vein, which drains the gastrointestinal tract. Briefly, after laparotomy, the portal vein was localized and the blood was collected with a syringe. Portal vein blood was processed following the same protocol above for peripheral blood.

### Evaluation of villi and crypt morphology in the small intestine

Small intestine samples (14 cm distal from the pylorus, 10cm long) were gently swiss rolled from the distal end and fixed in 4% paraformaldehyde (Electron Microscopy Science, Hatfield USA). Formalin-fixed tissues were then processed for paraffin embedding, cut into 5-micron thick sections and stained with hematoxylin and eosin (H&E) or Ki-67/hematoxylin as per protocol by the Experimental Pathology Core Laboratory at New York University (NY, USA). Measurement of villi and crypt length on H&E slides was performed by two independent researches in a blinded fashion using coded slides. Per sample, we analyzed a total of 6 slides with a distance of 100uM between each cut. For each slide, 10 villi/crypts were measured, as a total of 60/sample. The mean values of villi and crypt length were plotted. The imaging data were processed and analyzed using Image J software (NIH, Bethesda, MD). Quantification of frequency of Ki-67+ cells in the whole crypt region was performed in a blinded fashion using coded slides and the tools of Qupath software^41^.

### Tissue processing for ELISA

Distal Ileum (6 cm from the ileal-cecal junction) was collected and extensively washed to clean out fecal matter. The samples were weighed and, using a tissue homogenizer, extracted in PBS Tween 0.1% (with protease inhibitor) and centrifuged to remove tissue debris. The supernatant was frozen until measurement of VIP concentrations in the tissue.

### ELISA for Vasoactive Intestinal Peptide

VIP content was measured in the blood plasma or homogenized tissue following manufacturer’s recommendations (EIAM-VIP-1, RayBiotech).

### Measurement of plasma concentration of triglycerides

Peripheral blood was collected as described above and plasma was used to quantify triglyceride concentrations following manufacturer’s recommendations (Sigma-Aldrich, MAK266).

### Scanning Electron Microscopy

Scanning Electron Microscopy was performed on 1-1.5 cm pieces from terminal ileum (2cm above the ileal-cecal junction). Intestine was cut open and washed to remove fecal matter, pinned in dental wax and fixed for 2h with a 0.1M sodium cacodylate buffer (CB, pH 7.4) containing 2.5% glutaraldehyde and 2% paraformaldehyde. Samples were post fixed in 1% OsO4 for 2 hours, dehydrated in ethanol, and critical point dried using Tousimis autosamdsri 931 (Rockwille, MD). The dried intestines were put on SEM stabs, sputter coated with gold/palladium by DESK V TSC HP Denton Vacuum (Moorestown, NJ), and images were taken on random locations in the tissue by Zeiss Gemini300 FESEM using secondary electron mode at 5kv. For quantification of SFB length, random fields were selected for measurement using Image J.

### RNA extraction from intestinal epithelial cells, RT-PCR and qPCR

RNA isolation of CCR6^+^ ILC3, CCR6^neg^ ILC3, γδT17, Th17 and ileal epithelial cells, was performed using TRIzol following manufacter’s instructions (Invitrogen) followed by DNase I (Qiagen) treatment and cleanup with RNeasy MinElute kit (Qiagen) following manufacturer protocols. cDNA was generated using SuperScriptTM IV First-Strand Synthesis System (ThermoFisher). Gene-specific primers spanning exons were used: *Rps17* (F:’5-cgccattatccccagcaag-3’/ R:’5-tgtcgggatccacctcaatg-3’), *RegIIIγ* (F:’5-tctgcaagacagacaagatgct-3’/ R:’5-ggggcatctttcttggcaac-3’), *Fabp2* (F:’5-gtctagcagacggaacggag-3’/R:’5-ctccttcatatgtgtaggtctgga-3’), *Cd36* (F:’5-tggccttacttgggattgg-3’/R:’5-ccagtgtatatgtaggctcatcca-3’). Quantitative PCR was performed using the Hot Start-IT SYBRGreen (Affymetrix) on the Roche real-time PCR system (Roche 480). Values were normalized to *Rps17* gene for each sample. For evaluation of *Vipr2* mRNA expression, TaqMan probes *for Vipr2* (Mm01238618) and *Hprt* (Mm00446968) were purchased from Thermo Fisher Scientific and reactions were run with the CFX Connect Real—Time PCR Deterction System (Bio-Rad).

### Analysis of microbiota composition in the ileal fecal material, fecal pellet and ileal tissue

The ileum (ileal tissue: 1 cm), ileal fecal content, fecal pellet were collected from mice, and bacterial DNA was extracted with the QIAamp DNA Stool Kit (QIAGEN) following the manufacturer’s instructions. In the ileal tissue, SFB was quantified by qPCR with primers specific for 16S SFB rRNA genes (SFB F- sequence/SFB R- sequence)using Hot Start-IT SYBRGreen (Affymetrix) on the Roche real-time PCR system (Roche 480). The relative values of SFB in the tissue were normalized based on the amounts of host gDNA using primers specific for the genomic *Hif1a* locus (Hif1a F-sequence / Hif1a R- sequence). In the ileal fecal content and fecal pellet, SFB was quantified by qPCR with primers specific for SFB and the relative values were normalized based on the total relative amounts of 16S rDNA (16S F-sequence / 16S R-sequence).

For analysis of the whole microbial community composition in the ileal and fecal matter, sequencing of the 16S rRNA was performed as previously described^42^ in a MiSeq instrument (Ilumina, San Diego, USA) using 150bp, paired ended chemistry. 16S rRNA sequence data were processed using QIIME v2^43^.

### Data processing of publicly available RNA-seq

DESeq2-normalized gene quantification and differential expression analysis were downloaded from GSE116092^16^. Raw counts were downloaded from GSE127267 (ImmGen ULI RNA-seq data)^17^ and differential expression analysis was performed using DESeq2. Normalized counts were used for downstream analysis. A cutoff was made based on the normalized counts of a known non-expressed gene (*Foxp3*) (GSE116092, cut-off ≤ 9 and for GSE127267, cutoff <25).. GO term analysis of GSE116092^16^ was done using g:Profiler. For heat maps, genes were considered differentially expressed with FDR < 0.01 and log2 fold change ≥ 2.

### Library preparation for RNA sequencing and transcriptomic analysis

RNA-seq libraries for CCR6^+^ ILC3 were prepared with the TruSeq Stranded Total RNA Library Prep Gold Kit (Illumina, 20020598). The sequencing was performed using Illumina NextSeq instrument (Ilumina, San Diego, USA) using 150bp, paired ended chemistry. RNA-seq libraries were prepared and sequenced by the Genome Technology Core at New York University School of Medicine. Fastq files were aligned to the mouse Ensemble genome GRCm38 with STAR v 2.6.1d. Read pairs were counted using featurecounts from the Subread package v 1.6.2, prior to normalization and differential expression analysis which were performed using DESeq2 (Wald test with Benjamini–Hochberg correction to determine the FDR). Genes were considered differentially expressed with FDR<0.05 and log2 fold change > 1. Gene pathways and functions were assessed using Ingenuity Pathway Analysis (Qiagen Bioinformatics).

### Statistical analysis

Unpaired two-sided *t-test*, paired two-sided *t-test*, one-way ANOVA with multiple comparisons with Bonferroni correction, two-way ANOVA with multiple comparisons and Bonferroni correction, Mann-Whitney test, Mantel Cox test (for survival curves), were performed to compare the results using GraphPad Software Version 8 (GraphPad Software). No samples were excluded from analysis. We treated less than 0.05 p value as significant: \**P* < 0.05, *\*\*P* < 0.01, *\*\*\*P* < 0.001, and *\*\*\*\*P* < 0.0001.

## EXTENDED DATA FIGURE LEGENDS

**Extended Data Figure 1. Enrichment of transcripts related to nervous system/neural functions and development in CCR6**^**+**^ **ILC3. a**, Volcano-plot of differentially expressed genes between CCR6^+^ ILC3 and CCR6^neg^ ILC3 isolated from the small intestine of C57BL/6 mice GSE116092^16^. Green: Neurotransmitter/neuropeptide receptors, Blue: genes related to nervous system development/axonal guidance and contact. **b**, Top 10 Gene-Ontology terms from a comparison between subtypes of ILC3 showing enrichment of transcripts related to neuron differentiation and generation in CCR6^+^ ILC3 when compared to CCR6^neg^ ILC3. Green: Neurotransmitter receptors, Blue: genes related to nervous system development/axonal guidance and contact. **c**, Vocano-plot of differentially expressed genes between CCR6^+^ ILC3 (enriched in cryptopatches and ILFs^44^) and NKp46^+^ ILC3 (low presence in CPs and ILFs^45^) (GSE127267^17^).

**Extended Data Figure 2. Cryptopatch-associated enteric neurons are also present in the large intestine (colon) lamina propria. a, b** Representative immunofluorescence images of lamina propria neuronal projections of enteric neurons in the large intestine of *Rorc(t)*^EGFP/+^ mice. **(a)** Cluster of ILC3 (GFP^+^ cells, green) in close proximity of neuronal projections (βIII-Tubulin, red) of the enteric neurons in the colon lamina propria. **(b)** Cluster of ILC3 (GFP^+^ cells, blue) in close proximity of neuronal projections (βIII-Tubulin, red) of Vasoactive Intestinal Peptide^+^ enteric neurons (green) in the colon lamina propria.

**Extended Data Figure 3.** Neurochemical code of the cryptopatch-associated enteric neurons in the small intestine lamina propria. a-c, Representative immunofluorescence images of different subtypes of lamina propria neuronal projections of enteric neurons in the small intestine of *Rorc(t)*^EGFP/+^ mice. (a) Substance P (green) is not present in neuronal projections (βIII-Tubulin, red) localized inside CPs/ILFs (cluster of GFP^+^ cells, blue) in the lamina propria. Representative of 15 CPs/ILFs analyzed in the small intestine of 4 different *Rorc(t)*^EGFP/+^ mice). No colocalization of substance P was observed in any of the cryptopatch associated neuronal fibers. (b) Tyrosine hydroxylase^+^ neurons (green) are in close proximity but are not represented among neuronal projections (βIII-Tubulin, red) localized inside CPs/ILFs (cluster of GFP^+^ cells, blue) in the lamina propria. Representative of 20 CPs/ILFs analyzed in the small intestine of 4 different *Rorc(t)*^EGFP/+^ mice. TH^+^ fibers are found in proximity to ILC3 clusters but never intercalate with CPs/ILFs. (c) Vasoactive Intestinal Peptide^+^ (green) neurons (βIII-Tubulin, red) are in close proximity and interacting with ILC3 (GFP^+^, blue) in CPs/ILFs. Representative of 40 CPs/ILFs analyzed in the small intestine of 4 different *Rorc(t)*^EGFP/+^ mice.

**Extended Data Figure 4. VIP agonist inhibits *in vitro* IL-22 production by CCR6**^**+**^**ILC3. a**, *Vipr2* mRNA expression (relative to *Hprt*) in different subtypes of RORγt^+^ intestinal lymphoid cells, including ILC3 subsets, αβTh17 and γδT17 cells. RNA was isolated from CCR6^+^ ILC3, CCR6^negative^ ILC3, Th17 (αβ RORγt^+^T cells) and γδT17 (γδ RORγt^+^T cell) sorted from small intestine of *Rorc(t)-*Gfp^TG^ mice on basis of following markers: γdT17 cells (Lin^neg^RORgt^GFP+^CD3^+^TCRγ^+^), αβT17 (Lin^neg^RORγt^GFP+^CD3^+^TCRβ^+^), CCR6^+^ILC3 (Lin^neg^RORγt^GFP+^CD3^neg^ CCR6^+^) and CCR6^neg^ILC3 (Lin^neg^RORγt^GFP+^CD3^neg^CCR6^neg^) (n=4 mice). N.D= not detected. **b)** FACS plot showing gating strategy for identification and isolation of CCR6^+^ or CCR6^neg^ ILC3 (DAPI^neg^ Lin^neg^ CD127^+^ CD90.2^+^). **c, d**, *In vitro* activation of VIPR2 alone does not induce cytokine production or activation of CCR6^+^ ILC3. Representative FACS plots **(c)** and summary **(d)** for surface Sca-1 expression and intracellular IL-22 in small intestine lamina propria CCR6^+^ ILC3. N= 3, representative of 2 independent experiments. **e**, *In vitro* activation of VIPR2 promotes concentration-dependent inhibition of cytokine production by CCR6^+^ ILC3. Summary for IL-22 and IL-17a intracellular expression in small intestine lamina propria CCR6^+^ ILC3 stimulated *in vitro* for 12h with IL-23 (300pg/mL) with different concentrations of VIP. *\*P=< 0.05 ****P=< 0.001* (One-way ANOVA). Data are representative of two independent experiments. **f**, *In vitro* activation of VIPR2 does not reduce IL-23-induced Sca-1 expression on CCR6^+^ ILC3. Summary of Sca-1 expression in small intestine lamina propria CCR6^+^ ILC3 stimulated *in vitro* for 12h with IL-23 (100pg/mL) with/without the VIPR2 ligands BAY-559837 or VIP. N= 3/group, representative of 2 independent experiments. **g, h**, *In vitro* VIPR2 activation does not affect IL-23-induced IL-22 production by CCR6^neg^ ILC3. Representative FACS plots **(g)** and summaries **(h)** of surface Sca-1 expression and intracellular IL-22 in small intestine lamina propria CCR6^neg^ ILC3 stimulated *in vitro f*or 12h with IL-23 (100pg/mL) with/without combination with VIPR2 ligand BAY-559837 (1nM). N= 3, and data are representative of two independent experiments.

**Extended Data Figure 5. VIPR2 is required for *in vivo* inhibition of IL-22 production by CCR6**^**+**^**ILC3. a, b**, Mixed bone marrow chimeras, showing **(a)** no difference in frequency and ratio of WT (*Vipr2*^+/+^) vs VIPR2 KO (*Vipr2*^-/-^) CCR6^+^ILC3 in the ileum of mice receiving equal number of cells (N=17 mice, combined from 2 independent experiments) and **(b)** VIPR2-dependent inhibition of IL-22 production in WT (*Vipr2*^+/+^, CD45.1) versus VIPR2 KO (*Vipr2*^-/-^, CD45.2) CCR6^+^ILC3 in the ileum of chimeric mice. N=11, \*\*\*\**P*<0.0001 (*paired t-test*). Data are representative of two independent experiments. **c, d, e**, Transcriptomic profile showing differences between CCR6^+^ ILC3 among WT and VIPR2 KO isolated from mixed bone marrow chimeras. **(c)** Volcano plot and **(d)** heatmap of selected genes differentially expressed between CCR6^+^ ILC3 from *Vipr2*^+/+^ and *Vipr2*^-/-^ (FDR 5%, FC>1). **(e)** Analysis of pathways associated with genes upregulated in *Vipr2*^-/-^ CCR6^+^ ILC3 compared to *Vipr2*^+/+^ CCR6^+^ ILC3. **f, g**, Inactivation of *Vipr2* in ILC3 and T cells (*RORc(t* ^*Cre*^*Vipr2*^*fl/fl*^) does not affect **(f)** proportion or **(g)** number of CCR6^+^ ILC3 in the mouse ileum. *RORc(t)*^*Cre*^*Vipr2*^*+/+*^: N=8; *RORc(t)*^*Cre*^*Vipr2*^*fl/fl*^: N=6. **h, i** Inactivation of *Vipr2* (*RORc(t)*^*Cre*^*Vipr2*^*fl/fl*^) affects the frequency of IL-22 production in CCR6^+^ ILC3. Representative FACS plot **(h)** and summaries **(i)** indicating frequency of IL-22 production in CCR6^+^ ILC3 from the ileum of *RORc(t)*^*Cre*^*Vipr2*^*+/+*^(N=8) and *RORc(t)*^*Cre*^*Vipr2*^*fl/fl*^ (N=6) mice. \*\*\**P*<0.001 (*t-test*), representative of 3 independent experiments.**j, k**, Inactivation of *Vipr2* promotes a minor increase in the production of IL-17a in CCR6^+^ ILC3. Representative FACS plot **(j)** and summaries **(k)** indicating frequency of IL-22 production in CCR6^+^ ILC3 from the ileum of *RORc(t)*^*Cre*^*Vipr2*^*+/+*^(N=3) and *RORc(t)*^*Cre*^*Vipr2*^*fl/fl*^(N=3) mice. Representative of three independent experiments. **l, m**, Inactivation of *Vipr2* does not affect IL-17a or IL-22 production in CD3^+^RORγt^+^ lymphoid populations, namely CD3^+^ RORγt ^+^TCRγ^+^TCRβ^neg^ (γδT17) and CD3^+^ RORγt ^+^TCRγ^neg^TCRβ^+^ (Th17) cells. *RORc(t)*^*Cre*^*Vipr2*^*+/+*^(N=4) and *RORc(t)*^*Cre*^*Vipr2*^*fl/fl*^(N=4) mice. Representative of two independent experiments. FACS plot **(l)** and summaries **(m)** indicating frequency of IL-17 and IL22. N.S: not significant. **n, o**, Intestinal villi and crypt morphology (H&E) indicating the measurement of villi (red bracket) and crypt (green bracket) lengths. Values represent the average length of 60 structures (villi or crypt) per mouse, in a 10cm region, starting 15cm from the pylorus. \**P*<0.05 and \*\**P*<0.01 (*t-test*). *RORc(t)*^*Cre*^*Vipr2*^*+/+*^(N=4) and *RORc(t)*^*Cre*^*Vipr2*^*fl/fl*^ (N=3) mice. **p, q**, Ki-67 and Haematoxilin staining, revealing increased Ki-67^+^ cells in the crypt region of the small intestine (automated counting). \*\**P*<0.01 (*t-test*), *RORc(t)*^*Cre*^*Vipr2*^*+/+*^(N=4) and *RORc(t)*^*Cre*^*Vipr2*^*fl/fl*^(N=3) mice.

**Extended Data Figure 6. Effect of VIPergic neuronal modulation with DREADDs on IL-22 production by CCR6**^**+**^ **ILC3. a, b**, *VIP*^*Cre*^ activity in neurons in the gut. Homozygous *VIP* ^*Cre*^ mice were bred to homozygous *Rosa26*^*fl-st-fl-tdTomato*^. **(a)** Distribution of tdTomato^+^ projections in the small intestine lamina propia. tdTomato: red; nucleus/Dapi: blue. **(b)** tdTomato positive and negative neurons present in the small intestine. tdTomato: red; pan=neuronal marker β3-tubulin: green. Two distinct β3-tubulin^+^ tdTomato^+^ neuronal projections (white arrows) can be observed in a bundle of enteric neurons in the villi. Yellow: β3-tubulin^+^ tdTomato^negative^ projection. **c**, Nuclear cFOS localization in VIPergic neurons 2h after CNO (1mg/Kg, i.p.) or vehicle treatment of mice expressing the DREADD for activation under the control of *VIP*^*Cre*^ (VIP^Cre^ hM3Dq^fl/+^). cFOS-AF488: red, hM3Dq-TA-AF647: green, Nucleus/Dapi: blue. **d, e, f**, Effect of chemogenetic modulation of VIPergic neurons on IL-22 production by CCR6^+^ ILC3 in the small intestine (S.I.) and large intestine (L.I.). Representative FACS plot **(d)** and summaries of IL-22 production by CCR6^+^ ILC3 in mice expressing the DREADD for inhibition (hM4Di) **(e)** and for activation (hM3Dq) **(f)**. All the mice were treated with CNO (1mg/Kg, i.p, twice, 24h before sample collection). **(e):** N= 6 mice/group; **(f)** *VIP*^*Cre*^: N= 6, *VIP*^*Cre*^*hM3Dq*^*Cre*^: N= 5. \**P*<0.05, \*\*\*\**P*<0.001 (*t-test*). **g, h**, No difference in the frequencies of IL-17 and IL-22 production by CD3^+^ RORγt ^+^TCRγ^+^TCRβ^neg^ (γδT17) and CD3^+^ RORγt ^+^TCRγ^neg^TCRβ ^+^ (Th17) cells after activation of VIPergic neurons. Representative FACS plot **(g)** and summaries **(h).** All the mice were treated with CNO (1mg/Kg, i.p, twice, 24h before sample collection).

**Extended Data Figure 7. VIPergic neurons regulate host resistance to enteropathogenic *Citrobacter rodentium*. a**, Normalized *Vip* mRNA expression in the large intestine (cecum and proximal colon) of C57BL/6 mice at different time points after oral infection with *Citrobacter rodentium* (2×10^9^ CFU). **Day** 0: N=4; 2, 4 and 9 days after infection: N=6. \**P*<0.05 compared to day 0 (one-way ANOVA), representative of two independent experiments**. b, c**, Increased VIP activity in the gastrointestinal tract but not systemically in mice infected with *C. rodentium.* Concentrations of VIP in plasma from the (**b**) hepatic portal vein, which drains the gastrointestinal tract, and (**c**) peripheral blood of mice at different time points after intragastric administration of vehicle or *C. rodentium* (2×10^9^ CFU). d.p.i: days post-intragastric infection with *C. rodentium*. Data shown are pooled from two independent experiments. N= 8/group, *\*P*<0.05, *\*\*\*\*P*<0.0001(*t-test*). **d**, Off -target effects of CNO treatment do not account for mortality observed using activating DREADD during *C. rodentium* infection. Survival rates for *C. rodentium*-infected *Vip*^*IRES-Cre*^ *or Vip*^*IRES-Cre*^ *hM3Dq*^*fl-stop-fl/+*^ mice treated with CNO (1mg/Kg, daily, 1-4 d.p.i.: yellow rectangle). Vehicle: n=11, CNO: n=9. **e, f**, Infectious burden in feces of (**e**) *Vip*^*IRES-Cre*^*hM3Dq*^*fl-stop-fl/+*^(activating DREADD) and (**f**) *Vip*^*IRES-Cre*^*hM4Di* ^*fl-stop-fl/+*^(inhibitory DREADD) mice treated with vehicle or CNO (1mg/Kg, daily, 1-4 days post-intragastric infection with 2×10^9^ CFU for *Vip*^*IRES-Cre*^*hM3Dq*^*fl-stop-fl/+*^ and 4×10^10^ CFU for *Vip*^*IRES-Cre*^*hM4Di*^*fl-stop-fl/+*^mice). Log_10_ Colony Forming Units (CFU) of *C. rodentium* 9 days post-oral innoculation (9 d.p.i.). Data representative of two independent experiments. Activating DREADD mice: Vehicle: n=11, CNO: n=9. Inhibitory DREADD mice: Vehicle: n=8, CNO: n=7, \**P*=0.0009 (Mann-Whitney test). **g, h**, Exogenous treatment with recombinant murine IL-22 (rmIL-22, 250μg/mouse/day) protects against increase in **(g)** mortality and **(h)** bacterial dissemination to the spleen induced by VIPergic activation of *Vip*^*IRES-Cre*^*hM3Dq*^*fl-stop-fl/+*^mice. ***\*****P=0.0321 (Mantel Cox test*, survival); *\*\*P*=0.0022 (Mann-Whitney test). For visualization in the logarithm scale (e-g), CFU counts of 0 were attributed a value of 1. **i, j**, Inactivation of *Vipr2* expression in ILC3 (*RORc(t)*^*Cre*^*Vipr2*^*fl/fl*^) enhances barrier protection after oral infection with the enteropathogen *C. rodentium* (3×10^10^ CFU) **(i)** Discrete protection for weight loss in the first 9 days after infection. **(j**,) Log_10_ Colony Forming Units (CFU) of *C. rodentium* 9 days post-oral inoculation with 3×10^10^ CFU. *RORc(t)*^*Cre*^*Vipr2*^*fl/fl*^ mice display reduced amounts of *C. rodentium* translocation to the spleen and liver, and reduced CFU counts in the feces. *RORc(t)*^*Cre*^*Vipr2*^*+/+*^(N=10) and *RORc(t)*^*Cre*^*Vipr2*^*fl/fl*^ (N=10) mice. \**P*<0.05, \*\**P*<0.01 \*\*\**P*<0.001, \*\*\**P*<0.0001, [two-way ANOVA **(i)** and Mann-Whitney test **(j)**]. For visualization in the logarithm scale (e-g), CFU counts of 0 were attributed a value of 1.

**Extended Data Figure 8. Feeding controls intestinal VIP release and IL-22 production by CCR6**^**+**^ **ILC3 via activation of VIPR2. a**, Measurement of concentration of VIP in the ileum reveals higher amounts during dark-phase (feeding period, ZT12-ZT0) than in the light-phase (resting period, ZT0-ZT12). N=4, representative of two independent experiments. **b**, Concentrations of VIP in plasma isolated from hepatic portal vein blood of mice fed or fasted for 6 h. Blood samples were collected at two different time-points, during the light-phase period (ZT 6, 12PM) and the dark-phase period (ZT 18, 12AM). N=4, \**P*<0.05; \*\**P*<0.01 (*t-test*). **c**, Concentrations of VIP in plasma isolated from the peripheral blood of mice. Blood samples were collected at two time-points, during the light-phase period (ZT 6, 12PM) and the dark-phase period (ZT 18, 12AM). N=4, representative of two independent experiments. **d**, IL-22 expression by CCR6^+^ ILC3 from the ileum of CD45.1 *Vipr2*^+/+^:CD45.2 *Vipr2*^-/-^bone marrow chimeric mice at 6h after fasting (fasted, ZT 6) and 6h after feeding (fed, ZT 18). n=7/group, \*\**P*<0.01 (*paired t-test*). Data are representative of two pooled independent experiments. **e**, Ratio of IL-22-expressing cells, relative to Figure 3c, among CCR6^+^ ILC3 from the ileum of CD45.1 *Vipr2*^+/+^:CD45.2 *Vipr2*^-/-^bone marrow chimeric mice at 6h after fasting (Fasted, ZT 6) and 6h after feeding (Fed, ZT 18). n=7, \*\*\**P<*0.001 (*t-test*).

**Extended Data Figure 9. Feeding/fasting modulates growth of epithelium-associated segmented filamentous bacteria (SFB) and composition of fecal commensal microbiota. a**, Representative SEM images (magnifications: 1K, upper pannel, and 3K, lower pannel) showing epithelial-attached SFB in the ileum of mice 12 h after feeding (long filaments) or fasting (short-filaments, “stubbles”) at ZT 0. **b**, SFB lengths at different time points during the day in mice that had been fed for two weeks during the light-phase (Light-phase fed: ZT0 to ZT12, red) or during the dark-phase (dark-phase fed: ZT12 to ZT0, blue). **c**, Relative amounts (log) of SFB 16S measured in the ileal tissue by qPCR, normalized based on host genomic DNA quantity. Mice were fed for two weeks during the light-phase or during the dark-phase and the ileal tissue was collected at two different time points, ZT0 or ZT12. **d**, Weighted Unifrac Principal coordinate analysis of 16S rRNA composition in the fecal pellet of mice that had been fed for two weeks during the light-phase (circles) or during the dark-phase (diamonds). Fecal samples were collected from the same mice at different time points (ZT0 or ZT12). Dark-phase fed: at ZT0 these mice had been fed for 12h (red diamond), while at ZT12 they had fasted for 12h (blue diamond); Light-phase fed: at ZT0 these mice had fasted for 12h (blue circle), while at ZT12 they had been fed for 12h (red circle). **e**, Phylogenetic profile of the fecal microbiota composition associated with feeding/fasting status in the fecal pellet of mice that had been fed for two weeks during the light-phase or during the dark-phase.

**Extended Data Figure 10. Loss of *Vipr2* expression in ILC3 (*RORc(t)***^***Cre***^***Vipr2***^***fl/fl***^ **mice) affects the growth of epithelium-associated SFB and the composition of ileal and fecal commensal microbiota. a**, Relative amounts (log) of SFB 16S measured in the ileal tissue from *RORc(t)*^*Cre*^ *Vipr2*^*+/+*^or *RORc(t)*^*Cre*^ *Vipr2*^*fl/fl*^ mice (male, 8 weeks old) that had been fed for two weeks during the dark-phase. Samples were collected at ZT0 (12h fed). qPCR, normalized based on host gDNA levels. This cohort is composed of 3 different pairs of *RORc(t)*^*Cre*^ *Vipr2*^*+/+*^ and *RORc(t)*^*Cre*^ *Vipr2*^*fl/fl*^ littermates, housed in 3 different cages. **b**, Weighted Unifrac Principal coordinate analysis (PCoA) of 16S rRNA composition in the ileal fecal material from *RORc(t)*^*Cre*^*Vipr2*^*+/+*^ or *RORc(t)*^*Cre*^ *Vipr2*^*fl/fl*^ mice (male, 8 weeks old) that had been fed for two weeks during the dark-phase. Samples were collected at ZT0 (12h fed). This cohort was composed of 3 pairs of *RORc(t)*^*Cre*^ *Vipr2*^*+/+*^ and *RORc(t)*^*Cre*^ *Vipr2*^*fl/fl*^ littermates, housed in 3 different cages. **c**, Phylogenetic profile of the microbiota composition in the ileal fecal material of *RORc(t)*^*Cre*^ *Vipr22*^*+/+*^ or *RORc(t)*^*Cre*^ *Vipr2*^*fl/fl*^ mice after 12h of feeding. **d**, Weighted Unifrac Principal coordinate analysis (PCoA) of 16S rRNA composition in fecal pellet of the same mice described above from (*RORc(t)*^*Cre*^ *Vipr2*^*+/+*^ or *RORc(t)*^*Cre*^ *Vipr2*^*fl/fl*^ mice). **e**, Phylogenetic profile of the microbiota composition in the fecal pellet of *RORc(t)*^*Cre*^ *Vipr2*^*+/+*^ or *RORc(t)*^*Cre*^ *Vipr2*^*fl/fl*^ mice after 12h of feeding.

**Extended Data Figure 11. Feeding increase the efficiency of triglyceride absorption. a, b**, Plasma ^3^H CPM (counts per minute) in mice fed or fasted for 12h during the light-phase (ZT 0 – ZT 12, red and green circles) or during the dark-phase (ZT 12 – ZT 0, blue and black circles) and then gavaged with ^3^H-triolein and sampled at different times (**a**); and AUC during 4h **(b)**. AUC: Area under the curve per mL of plasma. N=4 mice per group, representative of two independent experiments, \**P* < 0.05 and \*\*\*\**P* < 0.001 (two-way ANOVA). **c**, Weight of 10 littermate/cagemate pairs of *RORc(t)*^*Cre*^ *Vipr2*^*+/+*^ and *RORc(t)*^*Cre*^ *Vipr2*^*fl/fl*^ mice under regular chow diet (male, 9 weeks old). Representative of two independent experiments, \*\**P* < 0.01 (*t-test*). **d**, Weight difference (%) between each *RORc(t)*^*Cre*^ *Vipr2*^*fl/fl*^ mouse compared to its paired *RORc(t)*^*Cre*^ *Vipr2*^*+/+*^ littermate/cagemate. **e**,**f**, Graphical abstract **(e)** and theoretical model **(f)** of the observations described created with BioRender.

## Supplementary Videos

**Supplementary Video 1.** 3D software reconstruction of Figure 1b showing the small intestine of *Rorc(t)*^EGFP/+^ mice with a cluster of intestinal ILC3 (cryptopatch) in close proximity to enteric neurons. Pan-neuronal marker: β3-tubulin^+^ (red), ILC3: GFP^+^ (green, respectively).

**Supplementary Video 2.** 3D software reconstruction of Figure 1c from the small intestine of *Rorc(t)*^EGFP/+^ mice showing ILC3 in the cryptopatch in close proximity to enteric neurons in the small intestine lamina propria. Pan-neuronal marker: β3-tubulin^+^ (red), ILC3: GFP^+^TCRβ^neg^ (green, respectively).

## REFERENCES

1. Constantinides MG. Interactions between the microbiota and innate and innate-like lymphocytes. J Leukoc Biol. 2018;103(3):409–419.

2. Sonnenberg GF. Regulation of intestinal health and disease by innate lymphoid cells. Int Immunol. 2014;26(9):501–507.

3. Spits H, Cupedo T. Innate lymphoid cells: emerging insights in development, lineage relationships, and function. Annu Rev Immunol. 2012;30:647–675.

4. Chayvialle JA, Miyata M, Rayford PL, Thompson JC. Effects of test meal, intragastric nutrients, and intraduodenal bile on plasma concentrations of immunoreactive somatostatin and vasoactive intestinal peptide in dogs. Gastroenterology. 1980;79(5 Pt 1):844–852.

5. Sano T, Huang W, Hall JA, et al. An IL-23R/IL-22 Circuit Regulates Epithelial Serum Amyloid A to Promote Local Effector Th17 Responses. Cell. 2015;163(2):381–393.

6. Savage AK, Liang HE, Locksley RM. The Development of Steady-State Activation Hubs between Adult LTi ILC3s and Primed Macrophages in Small Intestine. J Immunol. 2017;199(5):1912–1922.

7. Longman RS, Diehl GE, Victorio DA, et al. CX(3)CR1(+) mononuclear phagocytes support colitis-associated innate lymphoid cell production of IL-22. J Exp Med. 2014;211(8):1571–1583.

8. Mao K, Baptista AP, Tamoutounour S, et al. Innate and adaptive lymphocytes sequentially shape the gut microbiota and lipid metabolism. Nature. 2018;554(7691):255–259.

9. Zheng Y, Valdez PA, Danilenko DM, et al. Interleukin-22 mediates early host defense against attaching and effacing bacterial pathogens. Nat Med. 2008;14(3):282–289.

10. Ibiza S, Garcia-Cassani B, Ribeiro H, et al. Glial-cell-derived neuroregulators control type 3 innate lymphoid cells and gut defence. Nature. 2016;535(7612):440–443.

11. Mortha A, Chudnovskiy A, Hashimoto D, et al. Microbiota-dependent crosstalk between macrophages and ILC3 promotes intestinal homeostasis. Science. 2014;343(6178):1249288.

12. Hooper LV, Littman DR, Macpherson AJ. Interactions between the microbiota and the immune system. Science. 2012;336(6086):1268–1273.

13. Dudakov JA, Hanash AM, van den Brink MR. Interleukin-22: immunobiology and pathology. Annu Rev Immunol. 2015;33:747–785.

14. Eberl G, Marmon S, Sunshine MJ, Rennert PD, Choi Y, Littman DR. An essential function for the nuclear receptor RORgamma(t) in the generation of fetal lymphoid tissue inducer cells. Nat Immunol. 2004;5(1):64–73.

15. Gury-BenAri M, Thaiss CA, Serafini N, et al. The Spectrum and Regulatory Landscape of Intestinal Innate Lymphoid Cells Are Shaped by the Microbiome. Cell. 2016;166(5):1231–1246 e1213.

16. Pokrovskii M, Hall JA, Ochayon DE, et al. Characterization of Transcriptional Regulatory Networks that Promote and Restrict Identities and Functions of Intestinal Innate Lymphoid Cells. Immunity. 2019.

17. Heng TS, Painter MW, Immunological Genome Project C. The Immunological Genome Project: networks of gene expression in immune cells. Nat Immunol. 2008;9(10):1091–1094.

18. Furness JB. The enteric nervous system and neurogastroenterology. Nat Rev Gastroenterol Hepatol. 2012;9(5):286–294.

19. Muller PA, Koscso B, Rajani GM, et al. Crosstalk between muscularis macrophages and enteric neurons regulates gastrointestinal motility. Cell. 2014;158(2):300–313.

20. Cardoso V, Chesne J, Ribeiro H, et al. Neuronal regulation of type 2 innate lymphoid cells via neuromedin U. Nature. 2017;549(7671):277–281.

21. Klose CSN, Mahlakoiv T, Moeller JB, et al. The neuropeptide neuromedin U stimulates innate lymphoid cells and type 2 inflammation. Nature. 2017;549(7671):282–286.

22. Margolis KG, Gershon MD. Enteric Neuronal Regulation of Intestinal Inflammation. Trends Neurosci. 2016;39(9):614–624.

23. Zwarycz B, Gracz AD, Rivera KR, et al. IL22 Inhibits Epithelial Stem Cell Expansion in an Ileal Organoid Model. Cell Mol Gastroenterol Hepatol. 2019;7(1):1–17.

24. Zha JM, Li HS, Lin Q, et al. Interleukin 22 Expands Transit-Amplifying Cells While Depleting Lgr5(+) Stem Cells via Inhibition of Wnt and Notch Signaling. Cell Mol Gastroenterol Hepatol. 2019;7(2):255–274.

25. Roth BL. DREADDs for Neuroscientists. Neuron. 2016;89(4):683–694.

26. Conlin VS, Wu X, Nguyen C, et al. Vasoactive intestinal peptide ameliorates intestinal barrier disruption associated with Citrobacter rodentium-induced colitis. Am J Physiol Gastrointest Liver Physiol. 2009;297(4):G735–750.

27. Kohsaka A, Laposky AD, Ramsey KM, et al. High-fat diet disrupts behavioral and molecular circadian rhythms in mice. Cell Metab. 2007;6(5):414–421.

28. Park O, Wang H, Weng H, et al. In vivo consequences of liver-specific interleukin-22 expression in mice: Implications for human liver disease progression. Hepatology. 2011;54(1):252–261.

29. Smith PM, Howitt MR, Panikov N, et al. The microbial metabolites, short-chain fatty acids, regulate colonic Treg cell homeostasis. Science. 2013;341(6145):569–573.

30. McVey Neufeld KA, Perez-Burgos A, Mao YK, Bienenstock J, Kunze WA. The gut microbiome restores intrinsic and extrinsic nerve function in germ-free mice accompanied by changes in calbindin. Neurogastroenterol Motil. 2015;27(5):627–636.

31. McVey Neufeld KA, Mao YK, Bienenstock J, Foster JA, Kunze WA. The microbiome is essential for normal gut intrinsic primary afferent neuron excitability in the mouse. Neurogastroenterol Motil. 2013;25(2):183–e188.

32. Chayvialle JA, Miyata M, Rayford PL, Thompson JC. Release of vasoactive intestinal peptide by distention of the proximal stomach in dogs. Gut. 1980;21(9):745–749.

33. Harmar AJ, Marston HM, Shen S, et al. The VPAC(2) receptor is essential for circadian function in the mouse suprachiasmatic nuclei. Cell. 2002;109(4):497–508.

34. Nussbaum JC, Van Dyken SJ, von Moltke J, et al. Type 2 innate lymphoid cells control eosinophil homeostasis. Nature. 2013;502(7470):245–248.

35. Godinho-Silva C, Domingues RG, Rendas M, et al. Light-entrained and brain-tuned circadian circuits regulate ILC3s and gut homeostasis. Nature. 2019;574(7777):254–258.

## Supplementary References

36. Eberl G, Littman DR. Thymic origin of intestinal alphabeta T cells revealed by fate mapping of RORgammat+ cells. Science. 2004;305(5681):248–251.

37. Lochner M, Peduto L, Cherrier M, et al. In vivo equilibrium of proinflammatory IL-17+ and regulatory IL-10+ Foxp3+ RORgamma t+ T cells. J Exp Med. 2008;205(6):1381–1393.

38. Wang H, Yang H, Shivalila CS, et al. One-step generation of mice carrying mutations in multiple genes by CRISPR/Cas-mediated genome engineering. Cell. 2013;153(4):910–918.

39. Yang H, Wang H, Jaenisch R. Generating genetically modified mice using CRISPR/Casmediated genome engineering. Nat Protoc. 2014;9(8):1956–1968.

40. Zhang F, Zarkada G, Han J, et al. Lacteal junction zippering protects against diet-induced obesity. Science. 2018;361(6402):599–603.

41. Bankhead P, Loughrey MB, Fernandez JA, et al. QuPath: Open source software for digital pathology image analysis. Sci Rep. 2017;7(1):16878.

42. Caporaso JG, Lauber CL, Walters WA, et al. Ultra-high-throughput microbial community analysis on the Illumina HiSeq and MiSeq platforms. ISME J. 2012;6(8):1621–1624.

43. Bolyen E, Rideout JR, Dillon MR, et al. Reproducible, interactive, scalable and extensible microbiome data science using QIIME 2. Nat Biotechnol. 2019;37(8):852–857.

44. Lugering A, Ross M, Sieker M, et al. CCR6 identifies lymphoid tissue inducer cells within cryptopatches. Clin Exp Immunol. 2010;160(3):440–449.

45. Reynders A, Yessaad N, Vu Manh TP, et al. Identity, regulation and in vivo function of gut NKp46+RORgammat+ and NKp46+RORgammat-lymphoid cells. EMBO J. 2011;30(14):2934–2947.

