## Extended Figure Data for "VIP-producing enteric neurons interact with innate lymphoid cells to regulate feeding-dependent intestinal epithelial barrier functions"

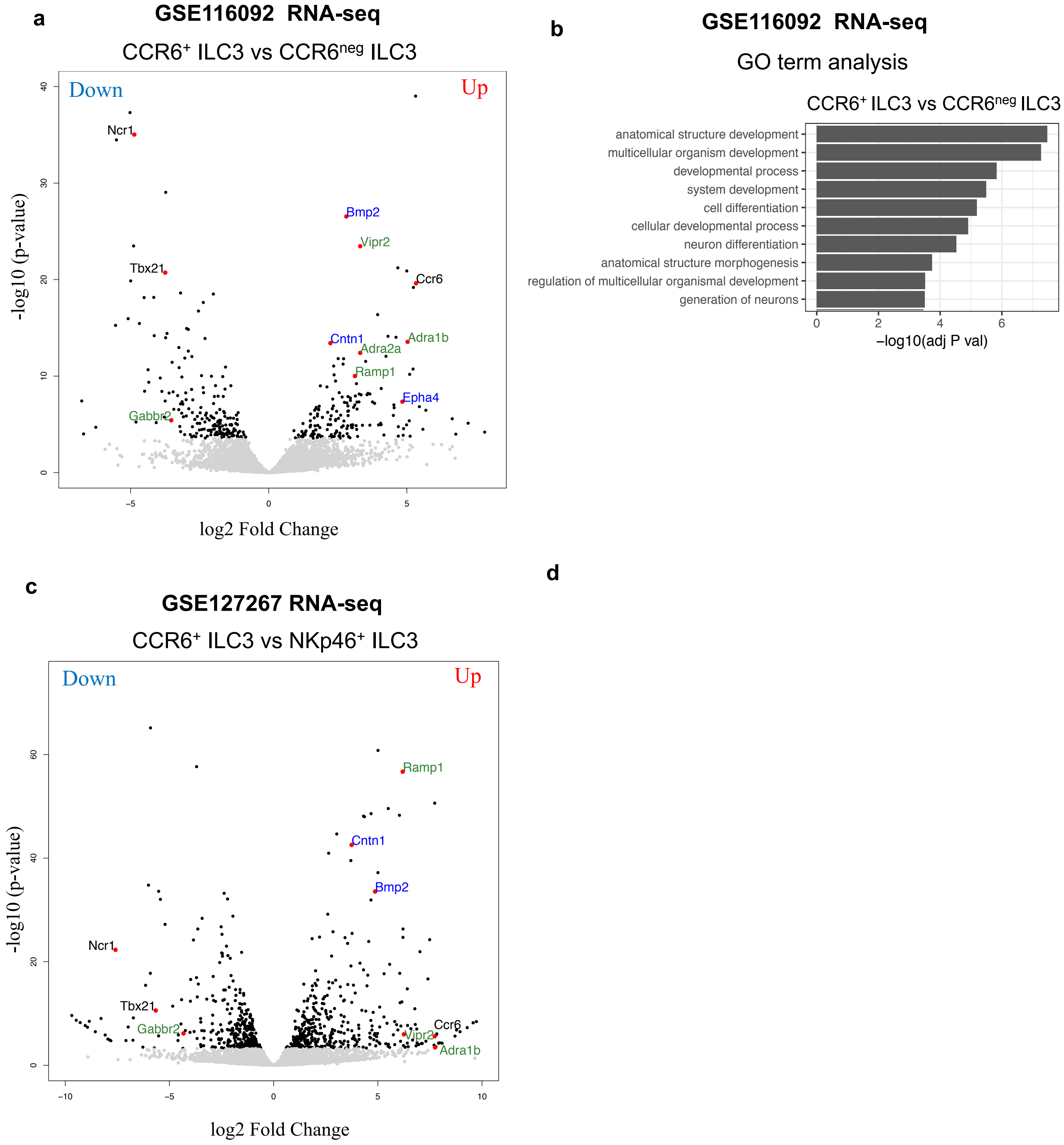


**Extended Data Figure 1. Enrichment of transcripts related to nervous system/neural functions and development in CCR6$^+$ ILC3. a,** Volcano-plot of differentially expressed genes between CCR6$^+$ ILC3 and CCR6$^{neg}$ ILC3 isolated from the small intestine of C57BL/6 mice GSE116092. Green: Neurotransmitter/ neuropeptide receptors, Blue: genes related to nervous system development/axonal guidance and contact. **b,** Top 10 Gene-Ontology terms from a comparison between subtypes of ILC3 showing enrichment of transcripts related to neuron differentiation and generation in CCR6$^+$ ILC3 when compared to CCR6$^{neg}$ ILC3. **c,** Vocano-plot of differentially expressed genes between CCR6$^+$ ILC3 (enriched in cryptopatches and ILFs) and NKp46$^+$ ILC3 (low presence in CPs and ILFs) (GSE127267). Green: Neurotransmitter receptors, Blue: genes related to nervous system development/axonal guidance and contact.

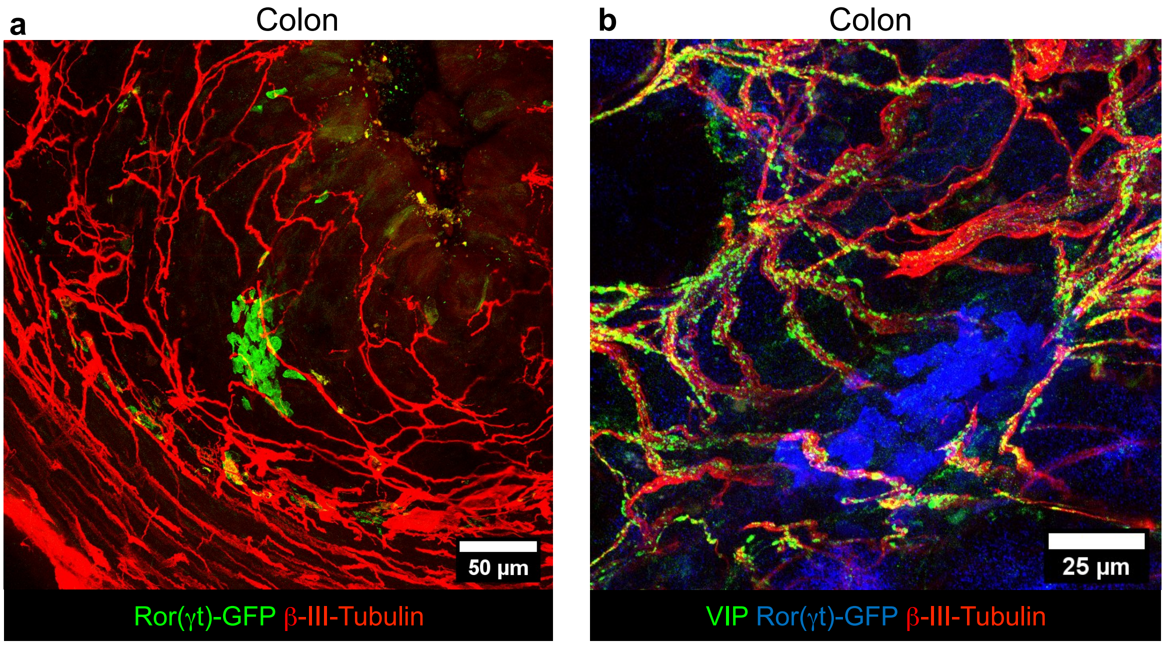


**Extended Data Figure 2. Cryptopatch-associated enteric neurons are also localized in the large intestine (colon) lamina propria. a, b** Representative immunofluorescence images of lamina propria neuronal projections of enteric neurons in the large intestine of *Rorc(γt)*$^{EGFP/+}$ mice. **(a)** Cluster of ILC3 (GFP$^{+}$ cells, green) in close proximity of neuronal projections (βIII-Tubulin, red) of the enteric neurons in the colon lamina propria. **(b)** Cluster of ILC3 (GFP$^{+}$ cells, blue) in close proximity of neuronal projections (βIII-Tubulin, red) of the Vasoactive Intestinal Peptide$^{+}$ enteric neurons (VIPen, green) in the colon lamina propria.

Extended Data Figure 3

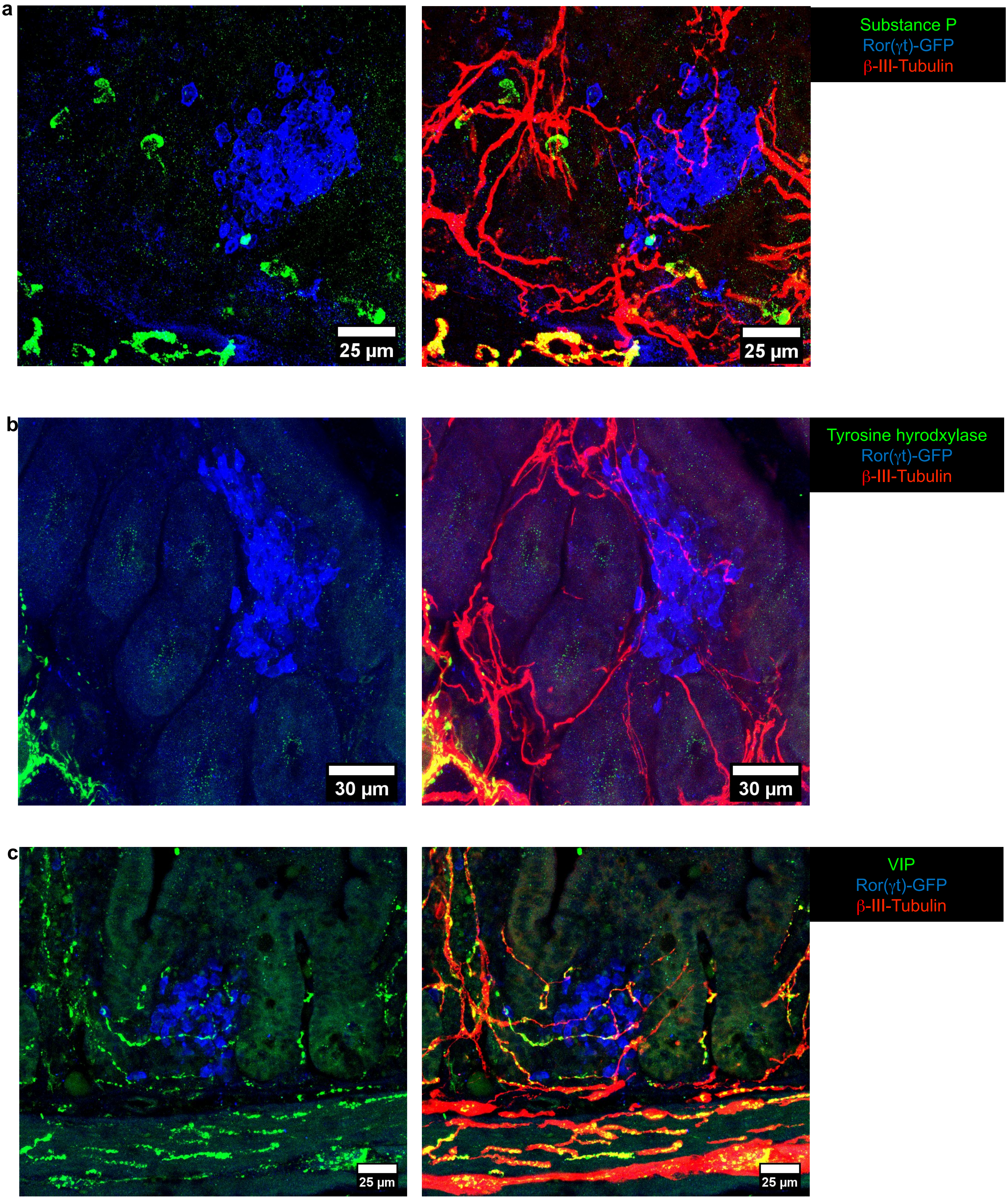

**Extended Data Figure 3.** Neurochemical code of the cryptopatch-associated enteric neurons in the small intestine lamina propria. a-c, Representative immunofluorescence images of different subtypes of lamina propria neuronal projections of enteric neurons in the small intestine of *Rorc(t)*$^{EGFP/+}$ mice. (a) Substance P (green) is not present in neuronal projections (βIII-Tubulin, red) localized inside CPs/ILFs (cluster of GFP$^{+}$ cells, blue) in the lamina propria. Representative of 15 CPs/ILFs analyzed in the small intestine of 4 different *Rorc(t)*$^{EGFP/+}$ mice). No colocalization of substance P was observed in any of the cryptopatch associated neuronal fibers. (b) Tyrosine hydroxylase$^{+}$ neurons (green) are in close proximity but are not represented among neuronal projections (βIII-Tubulin, red) localized inside CPs/ILFs (cluster of GFP$^{+}$ cells, blue) in the lamina propria. Representative of 20 CPs/ILFs analyzed in the small intestine of 4 different *Rorc(t)*$^{EGFP/+}$ mice. TH$^{+}$ fibers are found in proximity to ILC3 clusters but never intercalate with CPs/ILFs. (c) Vasoactive Intestinal Peptide$^{+}$ (green) neurons (βIII-Tubulin, red) are in close proximity and interacting with ILC3 (GFP$^{+}$, blue) in CPs/ILFs. Representative of 40 CPs/ILFs analyzed in the small intestine of 4 different *Rorc(t)*$^{EGFP/+}$ mice.

**Extended Data Figure 4**

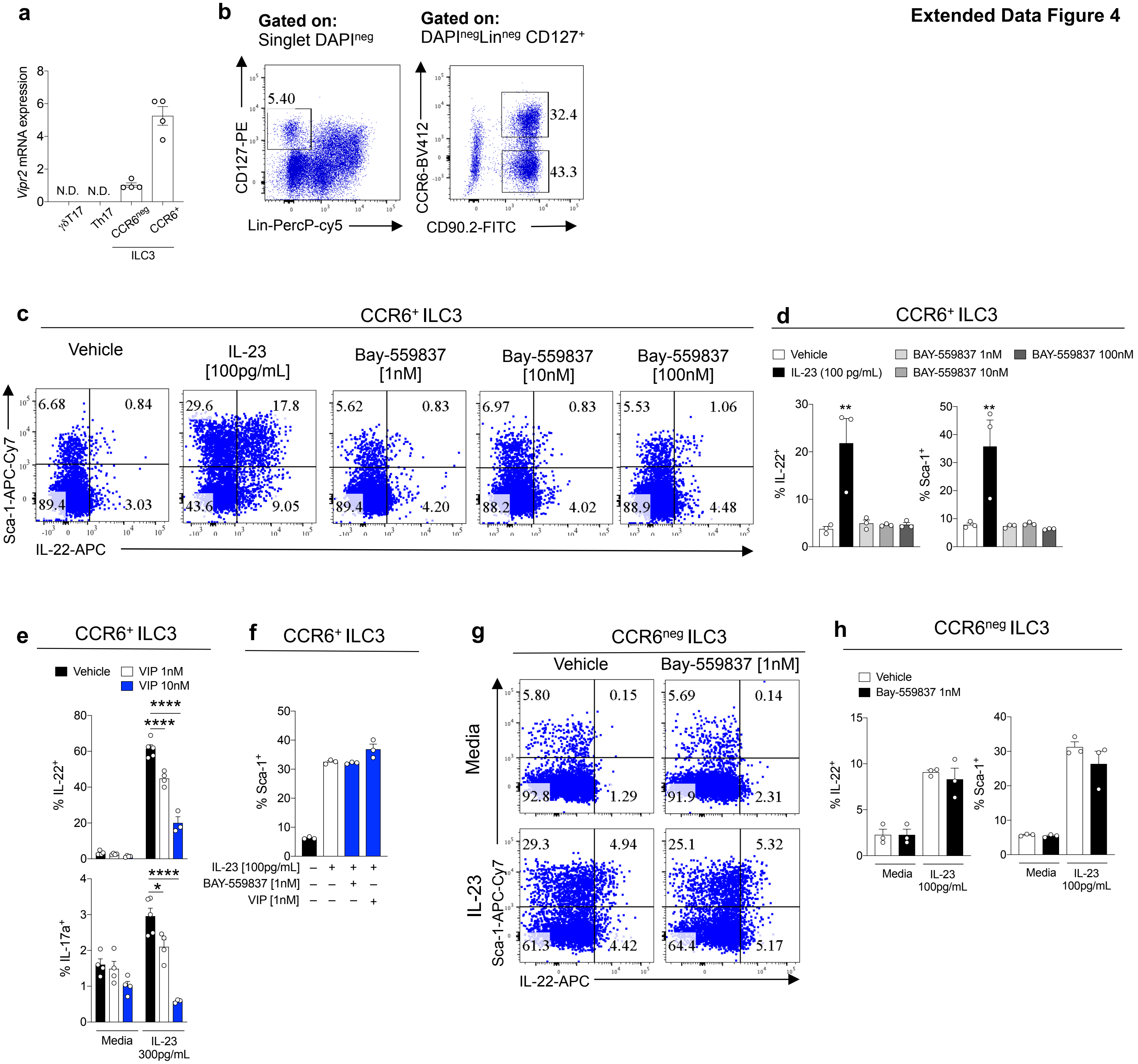

**Extended Data Figure 4. VIP agonist inhibits *in vitro* IL-22 production by CCR6$^{+}$ILC3. a,** *Vipr2* mRNA expression (relative to *Hprt*) in different subtypes of ROR$\gamma$t$^{+}$ intestinal lymphoid cells, including ILC3 subsets, $\alpha\beta$Th17 and $\gamma\delta$T17 cells. RNA was isolated from CCR6$^{+}$ ILC3, CCR6$^{negative}$ ILC3, Th17 ($\alpha\beta$ ROR$\gamma$t$^{+}$T cells) and $\gamma\delta$T17 ($\gamma\delta$ ROR$\gamma$t$^{+}$T cell) sorted from small intestine of *Rorc(t)*-Gfp$^{TG}$ mice on basis of following markers: $\gamma$dT17 cells (Lin$^{neg}$RORgt$^{GFP+}$CD3$^{+}$TCR$\gamma^{+}$), $\alpha\beta$T17 (Lin$^{neg}$ROR$\gamma$t$^{GFP+}$CD3$^{+}$TCR$\beta^{+}$), CCR6$^{+}$ILC3 (Lin$^{neg}$ROR$\gamma$t$^{GFP+}$CD3$^{neg}$ CCR6$^{+}$) and CCR6$^{neg}$ILC3 (Lin$^{neg}$ROR$\gamma$t$^{GFP+}$CD3$^{neg}$CCR6$^{neg}$) (n=4 mice). N.D= not detected. **b)** FACS plot showing gating strategy for identification and isolation of CCR6$^{+}$ or CCR6$^{neg}$ ILC3 (DAPI$^{neg}$Lin$^{neg}$ CD127$^{+}$ CD90.2$^{+}$). **c, d,** *In vitro* activation of VIPR2 alone does not induce cytokine production or activation of CCR6$^{+}$ ILC3. Representative FACS plots **(c)** and summary **(d)** for surface Sca-1 expression and intracellular IL-22 in small intestine lamina propria CCR6$^{+}$ ILC3. N= 3, representative of 2 independent experiments. **e,** *In vitro* activation of VIPR2 promotes concentration-dependent inhibition of cytokine production by CCR6$^{+}$ ILC3. Summary for IL-22 and IL-17a intracellular expression in small intestine lamina propria CCR6$^{+}$ ILC3 stimulated *in vitro* for 12h with IL-23 (300pg/mL) with different concentrations of VIP. **P=< 0.05 ****P=< 0.001* (One-way ANOVA). Data are representative of two independent experiments. **f,** *In vitro* activation of VIPR2 does not reduce IL-23-induced Sca-1 expression on CCR6$^{+}$ ILC3. Summary of Sca-1 expression in small intestine lamina propria CCR6$^{+}$ ILC3 stimulated *in vitro* for 12h with IL-23 (100pg/mL) with/without the VIPR2 ligands BAY-559837 or VIP. N= 3/group, representative of 2 independent experiments. **g, h,** *In vitro* VIPR2 activation does not affect IL-23-induced IL-22 production by CCR6$^{neg}$ ILC3. Representative FACS plots **(g)** and summaries **(h)** of surface Sca-1 expression and intracellular IL-22 in small intestine lamina propria CCR6$^{neg}$ ILC3 stimulated *in vitro f*or 12h with IL-23 (100pg/mL) with/without combination with VIPR2 ligand BAY-559837 (1nM). N= 3, and data are representative of two independent experiments.

### Extended Data Figure 5

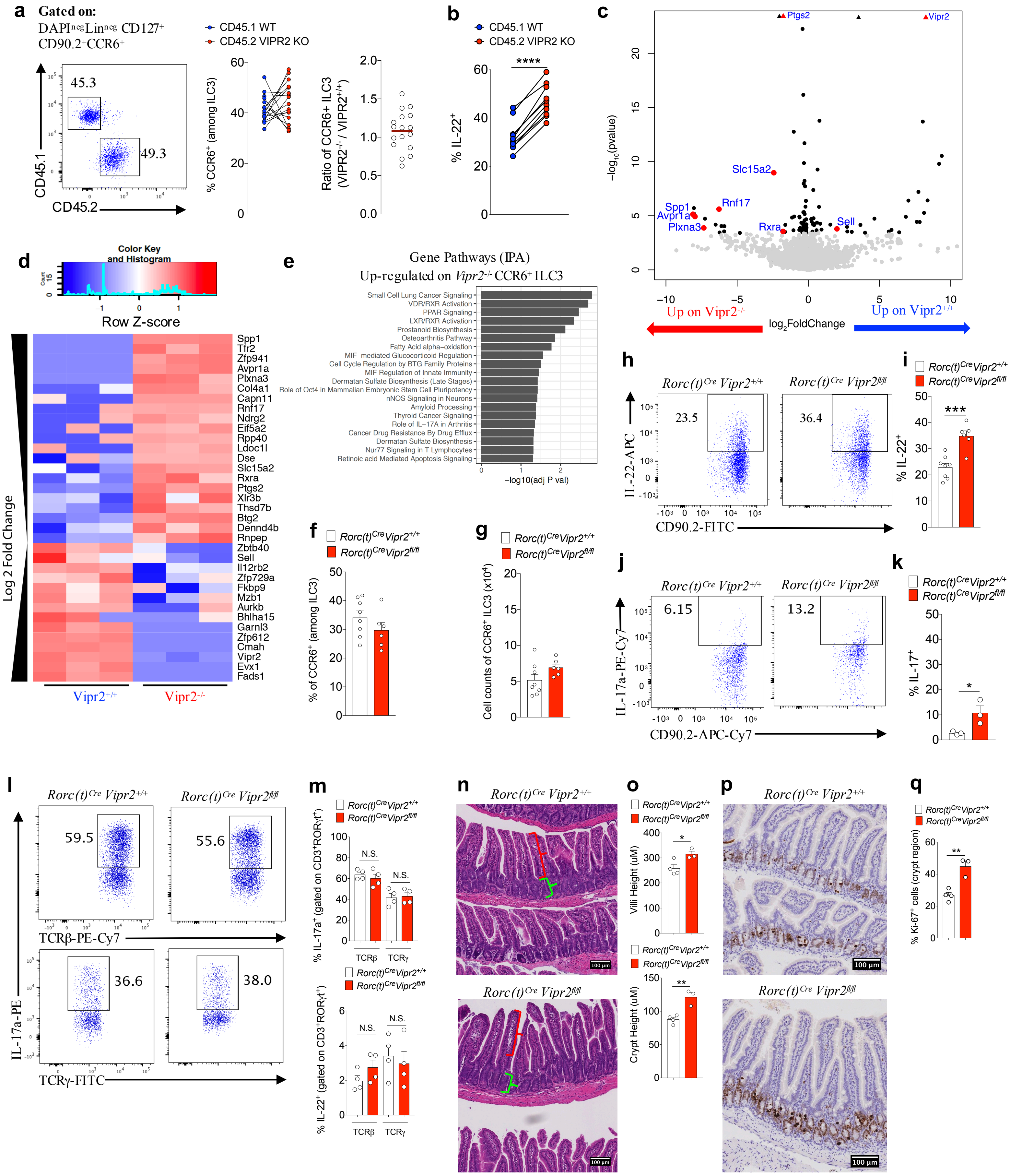

**Extended Data Figure 5. VIPR2 is required for *in vivo* inhibition of IL-22 production by CCR6$^+$ILC3. a, b,** Mixed bone marrow chimeras, showing **(a)** no difference in frequency and ratio of WT (*Vipr2*$^{+/+}$) vs VIPR2 KO (*Vipr2*$^{-/-}$) CCR6$^+$ILC3 in the ileum of mice receiving equal number of cells (N=17 mice, combined from 2 independent experiments) and **(b)** VIPR2-dependent inhibition of IL-22 production in WT (*Vipr2*$^{+/+}$, CD45.1) versus VIPR2 KO (*Vipr2*$^{-/-}$, CD45.2) CCR6$^+$ ILC3 in the ileum of chimeric mice. N=11, *****P*<0.0001 (*paired t-test*). Data are representative of two independent experiments. **c, d, e,** Transcriptomic profile showing differences between CCR6$^+$ ILC3 among WT and VIPR2 KO isolated from mixed bone marrow chimeras. **(c)** Volcano plot and **(d)** heatmap of selected genes differentially expressed between CCR6$^+$ ILC3 from *Vipr2*$^{+/+}$ and *Vipr2*$^{-/-}$ (FDR 5%, FC>1). **(e)** Analysis of pathways associated with genes upregulated in *Vipr2*$^{-/-}$ CCR6$^+$ ILC3 compared to *Vipr2*$^{+/+}$ CCR6$^+$ ILC3. **f, g,** Inactivation of *Vipr2* in ILC3 and T cells (*RORc(t)*$^{Cre}$*Vipr2*$^{fl/fl}$) does not affect **(f)** proportion or **(g)** number of CCR6$^+$ ILC3 in the mouse ileum. *RORc(t)*$^{Cre}$*Vipr2*$^{+/+}$: N=8; *RORc(t)*$^{Cre}$*Vipr2*$^{fl/fl}$: N=6. **h, i** Inactivation of *Vipr2* (*RORc(t)*$^{Cre}$*Vipr2*$^{fl/fl}$) affects the frequency of IL-22 production in CCR6$^+$ ILC3. Representative FACS plot **(h)** and summaries **(i)** indicating frequency of IL-22 production in CCR6$^+$ ILC3 from the ileum of *RORc(t)*$^{Cre}$*Vipr2*$^{+/+}$(N=8) and *RORc(t)*$^{Cre}$*Vipr2*$^{fl/fl}$ (N=6) mice. ****P*<0.001 (*t-test*), representative of 3 independent experiments.**j, k,** Inactivation of *Vipr2* promotes a minor increase in the production of IL-17a in CCR6$^+$ ILC3. Representative FACS plot **(j)** and summaries **(k)** indicating frequency of IL-22 production in CCR6$^+$ ILC3 from the ileum of *RORc(t)*$^{Cre}$*Vipr2*$^{+/+}$(N=3) and *RORc(t)*$^{Cre}$*Vipr2*$^{fl/fl}$ (N=3) mice. Representative of three independent experiments. **l, m,** Inactivation of *Vipr2* does not affect IL-17a or IL-22 production in CD3$^+$ROR$\gamma$t$^+$ lymphoid populations, namely CD3$^+$ ROR$\gamma$t $^+$TCR$\gamma^+$TCR$\beta^{neg}$ ($\gamma\delta$T17) and CD3$^+$ ROR$\gamma$t $^+$TCR$\gamma^{neg}$TCR$\beta^+$ (Th17) cells. *RORc(t)*$^{Cre}$*Vipr2*$^{+/+}$(N=4) and *RORc(t)*$^{Cre}$*Vipr2*$^{fl/fl}$ (N=4) mice. Representative of two independent experiments. FACS plot **(l)** and summaries **(m)** indicating frequency of IL-17 and IL22. N.S: not significant. **n, o,** Intestinal villi and crypt morphology (H&E) indicating the measurement of villi (red bracket) and crypt (green bracket) lengths. Values represent the average length of 60 structures (villi or crypt) per mouse, in a 10cm region, starting 15cm from the pylorus. **P*<0.05 and ***P*<0.01 (*t-test*). *RORc(t)*$^{Cre}$*Vipr2*$^{+/+}$(N=4) and *RORc(t)*$^{Cre}$*Vipr2*$^{fl/fl}$ (N=3) mice. **p, q,** Ki-67 and Haematoxilin staining, revealing increased Ki-67$^+$ cells in the crypt region of the small intestine (automated counting). ***P*<0.01 (*t-test*), *RORc(t)*$^{Cre}$*Vipr2*$^{+/+}$(N=4) and *RORc(t)*$^{Cre}$*Vipr2*$^{fl/fl}$ (N=3) mice.

**Extended Data Figure 6**

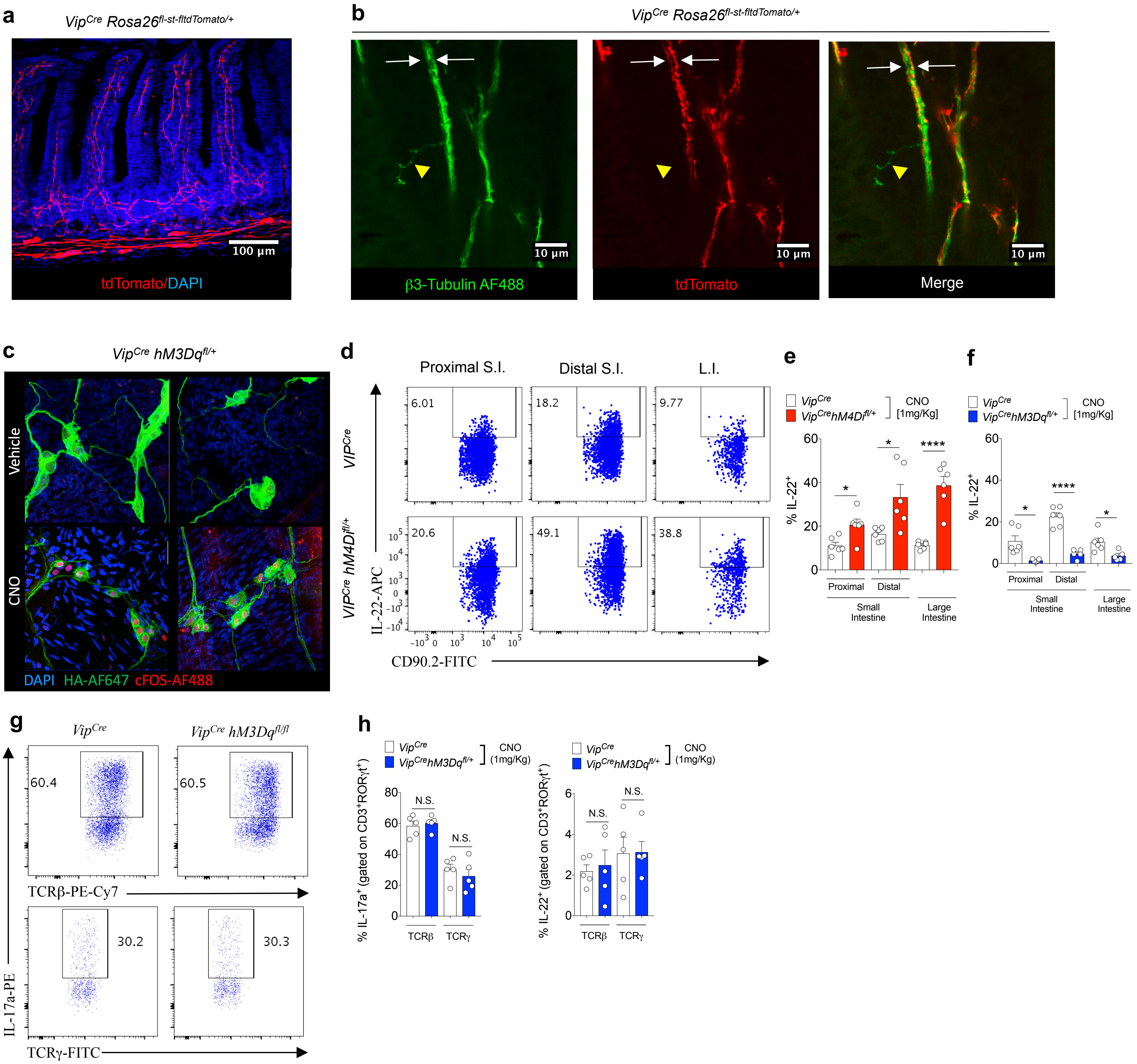


**Extended Data Figure 6. Effect of VIPergic neuronal modulation with DREADDs on IL-22 production by CCR6⁺ ILC3. a, b,** *VIP*$^{Cre}$ activity in neurons in the gut. Homozygous *VIP*$^{Cre}$ mice were bred to homozygous *Rosa26*$^{fl\text{-}st\text{-}fl\text{-}tdTomato}$. **(a)** Distribution of tdTomato$^{+}$ projections in the small intestine lamina propia. tdTomato: red; nucleus/Dapi: blue. **(b)** tdTomato positive and negative neurons present in the small intestine. tdTomato: red; pan=neuronal marker β3-tubulin: green. Two distinct β3-tubulin$^{+}$ tdTomato$^{+}$ neuronal projections (white arrows) can be observed in a bundle of enteric neurons in the villi. Yellow: β3-tubulin$^{+}$ tdTomato$^{negative}$ projection. **c,** Nuclear cFOS localization in VIPergic neurons 2h after CNO (1mg/Kg, i.p.) or vehicle treatment of mice expressing the DREADD for activation under the control of *VIP*$^{Cre}$ (VIP$^{Cre}$ hM3Dq$^{fl/+}$). cFOS-AF488: red, hM3Dq-TA-AF647: green, Nucleus/ Dapi: blue. **d, e, f,** Effect of chemogenetic modulation of VIPergic neurons on IL-22 production by CCR6$^{+}$ ILC3 in the small intestine (S.I.) and large intestine (L.I.). Representative FACS plot **(d)** and summaries of IL-22 production by CCR6$^{+}$ ILC3 in mice expressing the DREADD for inhibition (hM4Di) **(e)** and for activation (hM3Dq) **(f)**. All the mice were treated with CNO (1mg/Kg, i.p, twice, 24h before sample collection). **(e):** N= 6 mice/group; **(f)** *VIP*$^{Cre}$: N= 6, *VIP*$^{Cre}$*hM3Dq*$^{Cre}$ : N= 5. *$P<0.05$, ****$P<0.001$ (*t-test*). **g, h,** No difference in the frequencies of IL-17 and IL-22 production by CD3$^{+}$ RORγt$^{+}$TCRγ$^{+}$TCRβ$^{neg}$ (γδT17) and CD3$^{+}$ RORγt$^{+}$TCRγ$^{neg}$TCRβ$^{+}$ (Th17) cells after activation of VIPergic neurons. Representative FACS plot **(g)** and summaries **(h)**. All the mice were treated with CNO (1mg/Kg, i.p, twice, 24h before sample collection).

**Extended Data Figure 7**

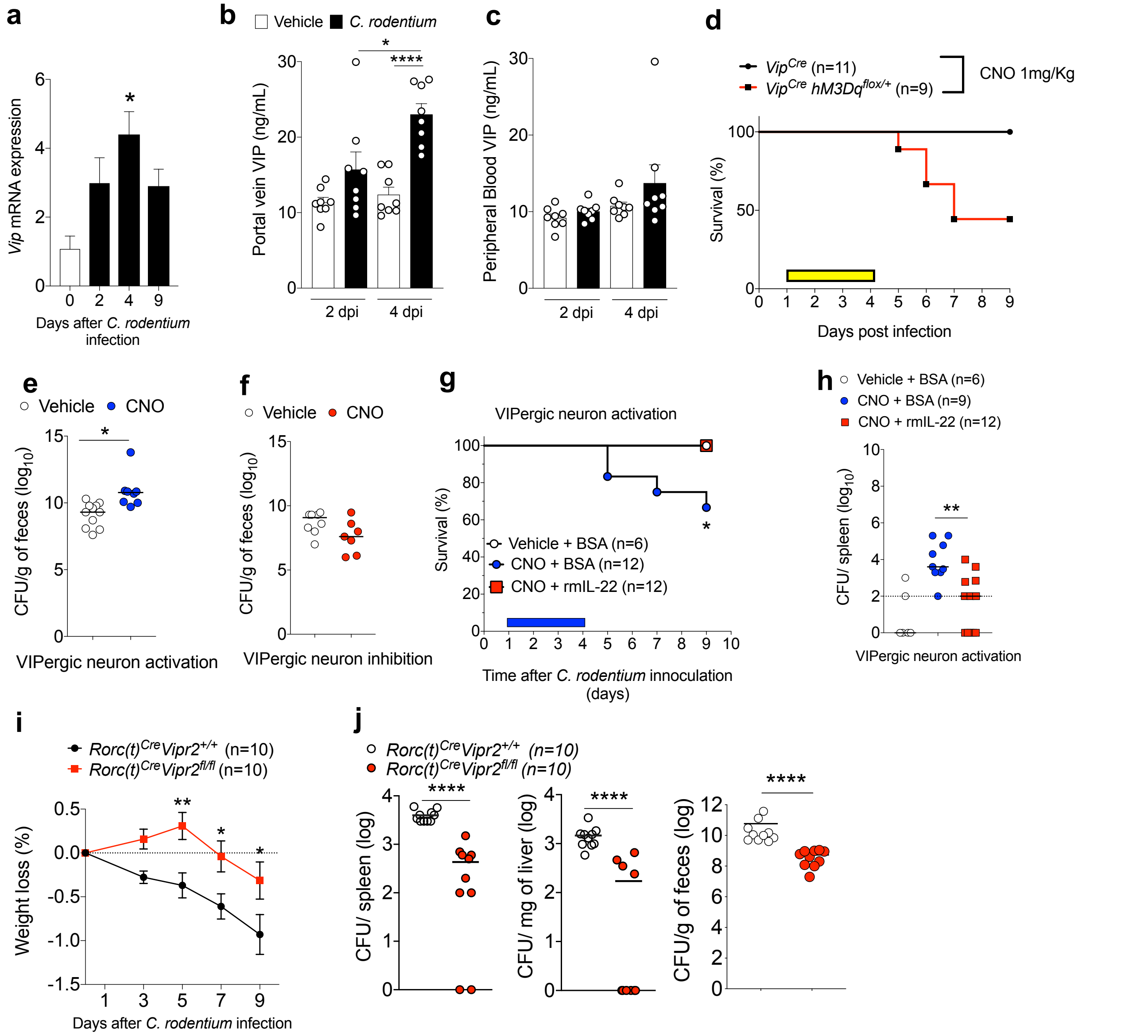


**Extended Data Figure 7. VIPergic neurons regulate host resistance to enteropathogenic *Citrobacter rodentium*. a,** Normalized *Vip* mRNA expression in the large intestine (cecum and proximal colon) of C57BL/6 mice at different time points after oral infection with *Citrobacter rodentium* (2x$10^9$ CFU). **Day** 0: N=4; 2, 4 and 9 days after infection: N=6. *$P$<0.05 compared to day 0 (one-way ANOVA), representative of two independent experiments**. b, c,** Increased VIP activity in the gastrointestinal tract but not systemically in mice infected with *C. rodentium.* Concentrations of VIP in plasma from the (**b**) hepatic portal vein, which drains the gastrointestinal tract, and (**c**) peripheral blood of mice at different time points after intragastric administration of vehicle or *C. rodentium* (2x$10^9$ CFU). d.p.i: days post-intragastric infection with *C. rodentium*. Data shown are pooled from two independent experiments. N= 8/group, *$P$<0.05, ****$P$<0.0001(*t-test*). **d,** Off -target effects of CNO treatment do not account for mortality observed using activating DREADD during *C. rodentium* infection**.** Survival rates for *C. rodentium*-infected $Vip^{IRES\text{-}Cre}$ *or* $Vip^{IRES\text{-}Cre}hM3Dq^{fl\text{-}stop\text{-}fl/+}$ mice treated with CNO (1mg/Kg, daily, 1-4 d.p.i.: yellow rectangle). Vehicle: n=11, CNO: n=9. **e, f,** Infectious burden in feces of (**e**) $Vip^{IRES\text{-}Cre}hM3Dq^{fl\text{-}stop\text{-}fl/+}$ (activating DREADD) and (**f**) $Vip^{IRES\text{-}Cre}hM4Di^{fl\text{-}stop\text{-}fl/+}$ (inhibitory DREADD) mice treated with vehicle or CNO (1mg/Kg, daily, 1-4 days post-intragastric infection with 2x$10^9$ CFU for $Vip^{IRES\text{-}Cre}hM3Dq^{fl\text{-}stop\text{-}fl/+}$ and 4x$10^{10}$ CFU for $Vip^{IRES\text{-}Cre}hM4Di^{fl\text{-}stop\text{-}fl/+}$ mice). $Log_{10}$ Colony Forming Units (CFU) of *C. rodentium* 9 days post-oral innoculation (9 d.p.i.). Data representative of two independent experiments. Activating DREADD mice: Vehicle: n=11, CNO: n=9. Inhibitory DREADD mice: Vehicle: n=8, CNO: n=7**,** *$P$=0.0009 (Mann-Whitney test). **g, h,** Exogenous treatment with recombinant murine IL-22 (rmIL-22, 250μg/mouse/day) protects against increase in **(g)** mortality and **(h)** bacterial dissemination to the spleen induced by VIPergic activation of $Vip^{IRES\text{-}Cre}hM3Dq^{fl\text{-}stop\text{-}fl/+}$mice. **$P$=0.0321 (Mantel Cox test,* survival); **$P$=0.0022 *(*Mann-Whitney test). For visualization in the logarithm scale (e-g), CFU counts of 0 were attributed a value of 1. **i, j,** Inactivation of *Vipr2* expression in ILC3 ($RORc(t)^{Cre}Vipr2^{fl/fl}$*)* enhances barrier protection after oral infection with the enteropathogen *C. rodentium* (3x$10^{10}$ CFU) **(i)** Discrete protection for weight loss in the first 9 days after infection. **(j,)** $Log_{10}$ Colony Forming Units (CFU) of *C. rodentium* 9 days post-oral inoculation with 3x$10^{10}$ CFU. $RORc(t)^{Cre}Vipr2^{fl/fl}$ mice display reduced amounts of *C. rodentium* translocation to the spleen and liver, and reduced CFU counts in the feces**.** $RORc(t)^{Cre}Vipr2^{+/+}$(N=10) and $RORc(t)^{Cre}Vipr2^{fl/fl}$ (N=10) mice. *$P$<0.05, **$P$<0.01 ***$P$<0.001, ***$P$<0.0001, [two-way ANOVA **(i)** and Mann-Whitney test **(j)**]. For visualization in the logarithm scale (e-g), CFU counts of 0 were attributed a value of 1.

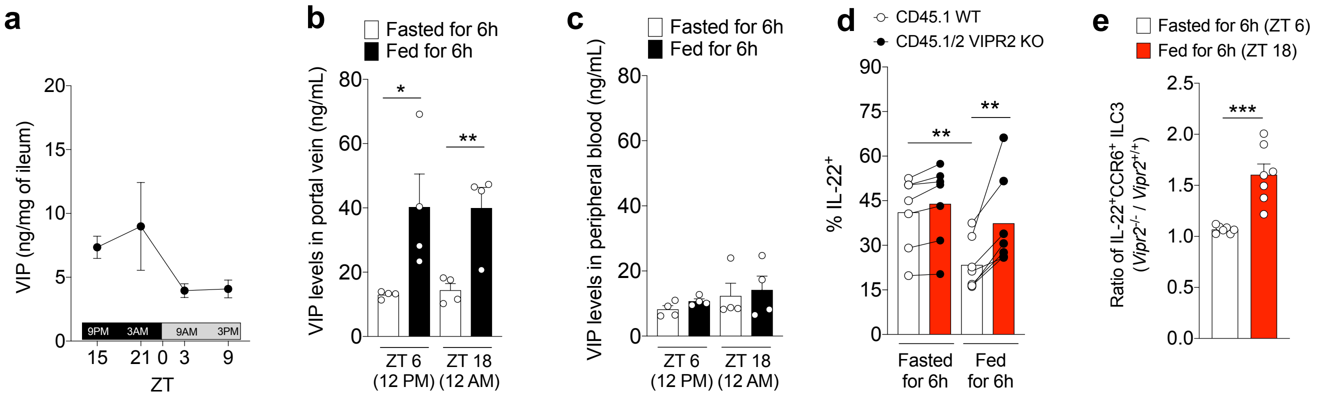


**Extended Data Figure 8. Feeding controls intestinal VIP release and IL-22 production by CCR6⁺ ILC3 via activation of VIPR2. a,** Measurement of concentration of VIP in the ileum reveals higher amounts during dark-phase (feeding period, ZT12-ZT0) than in the light-phase (resting period, ZT0-ZT12). N=4, representative of two independent experiments. **b,** Concentrations of VIP in plasma isolated from hepatic portal vein blood of mice fed or fasted for 6 h. Blood samples were collected at two different time-points, during the light-phase period (ZT 6, 12PM) and the dark-phase period (ZT 18, 12AM). N=4, **P*<0.05*;* ***P*<0.01 (*t-test*)*.* **c,** Concentrations of VIP in plasma isolated from the peripheral blood of mice. Blood samples were collected at two time-points, during the light-phase period (ZT 6, 12PM) and the dark-phase period (ZT 18, 12AM). N=4, representative of two independent experiments. **d,** IL-22 expression by CCR6⁺ ILC3 from the ileum of CD45.1 *Vipr2*$^{+/+}$:CD45.2 *Vipr2*$^{-/-}$ bone marrow chimeric mice at 6h after fasting (fasted, ZT 6) and 6h after feeding (fed, ZT 18). n=7/group, ***P*<0.01 (*paired t-test*). Data are representative of two pooled independent experiments. **e,** Ratio of IL-22-expressing cells, relative to Figure 3c, among CCR6⁺ ILC3 from the ileum of CD45.1 *Vipr2*$^{+/+}$:CD45.2 *Vipr2*$^{-/-}$ bone marrow chimeric mice at 6h after fasting (Fasted, ZT 6) and 6h after feeding (Fed, ZT 18). n=7, ****P*<0.001 (*t-test*).

**Extended Data Figure 9**

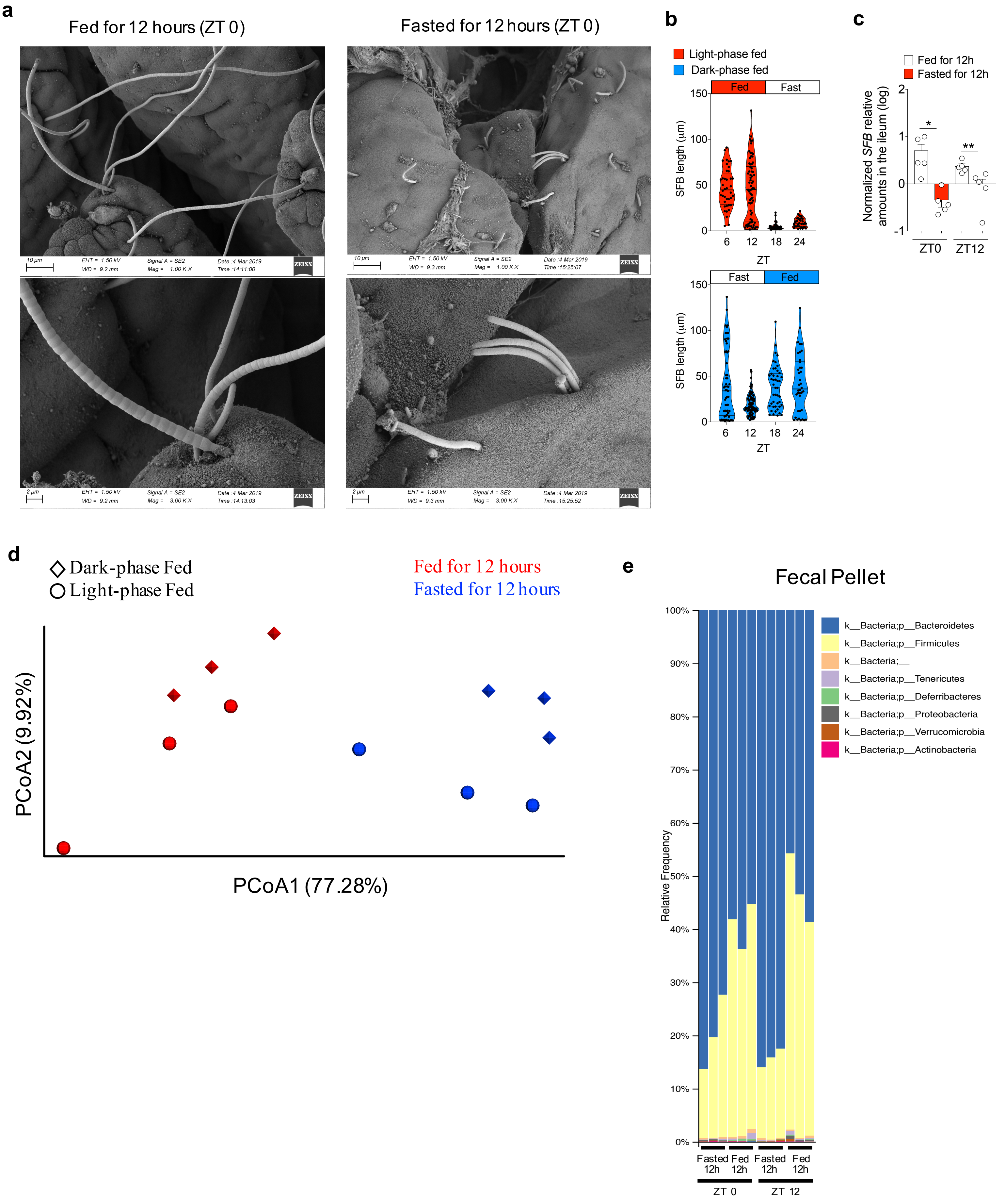

**Extended Data Figure 9. Feeding/fasting modulates growth of epithelium-associated segmented filamentous bacteria (SFB) and composition of fecal commensal microbiota. a,** Representative SEM images (magnifications: 1K, upper pannel, and 3K, lower pannel) showing epithelial-attached SFB in the ileum of mice 12 h after feeding (long filaments) or fasting (short-filaments, “stubbles”) at ZT 0. **b,** SFB lengths at different time points during the day in mice that had been fed for two weeks during the light-phase (Light-phase fed: ZT0 to ZT12, red) or during the dark-phase (dark-phase fed: ZT12 to ZT0, blue). **c,** Relative amounts (log) of SFB 16S measured in the ileal tissue by qPCR, normalized based on host genomic DNA quantity. Mice were fed for two weeks during the light-phase or during the dark-phase and the ileal tissue was collected at two different time points, ZT0 or ZT12. **d,** Weighted Unifrac Principal coordinate analysis (PCoA) of 16S rRNA composition in the fecal pellet of mice that had been fed for two weeks during the light-phase (circles) or during the dark-phase (diamonds). Fecal samples were collected from the same mice at different time points (ZT0 or ZT12). Dark-phase fed: at ZT0 these mice had been fed for 12h (red diamond), while at ZT12 they had fasted for 12h (blue diamond); Light-phase fed: at ZT0 these mice had fasted for 12h (blue circle), while at ZT12 they had been fed for 12h (red circle). **e,** Phylogenetic profile of the fecal microbiota composition associated with feeding/fasting status in the fecal pellet of mice that had been fed for two weeks during the light-phase or during the dark-phase.

Extended Data Figure 10

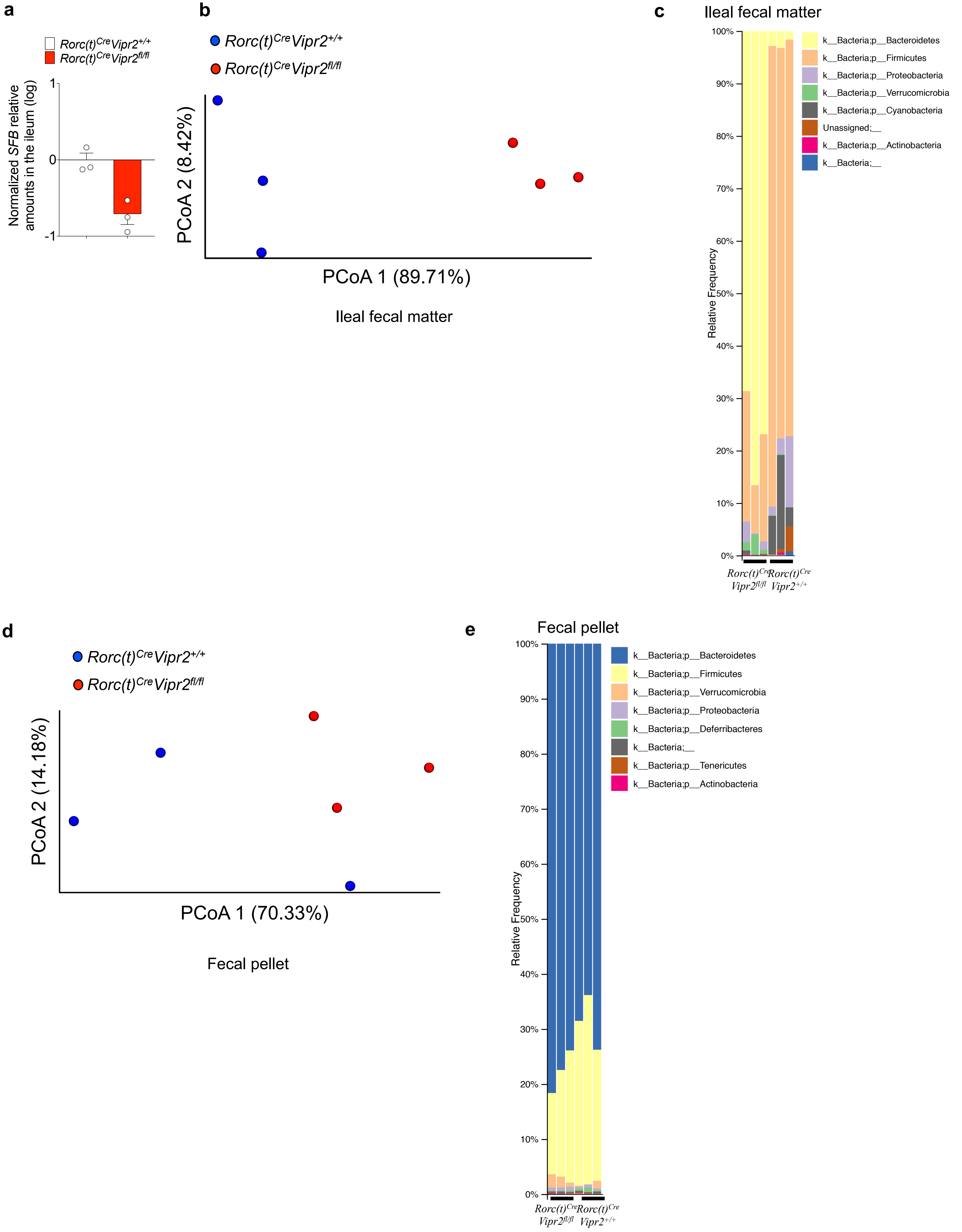

**Extended Data Figure 10. Loss of *Vipr2* expression in ILC3 (*RORc(t)$^{Cre}$Vipr2$^{fl/fl}$* mice) affects the growth of epithelium-associated SFB and the composition of ileal and fecal commensal microbiota. a,** Relative amounts (log) of SFB 16S measured in the ileal tissue from *RORc(t)$^{Cre}$ Vipr2$^{+/+}$* or *RORc(t)$^{Cre}$ Vipr2$^{fl/fl}$* mice (male, 8 weeks old) that had been fed for two weeks during the dark-phase. Samples were collected at ZT0 (12h fed). qPCR, normalized based on host gDNA levels. This cohort is composed of 3 different pairs of *RORc(t)$^{Cre}$ Vipr2$^{+/+}$* and *RORc(t)$^{Cre}$ Vipr2$^{fl/fl}$* littermates, housed in 3 different cages. **b,** Weighted Unifrac Principal coordinate analysis (PCoA) of 16S rRNA composition in the ileal fecal material from *RORc(t)$^{Cre}$ Vipr2$^{+/+}$* or *RORc(t)$^{Cre}$ Vipr2$^{fl/fl}$* mice (male, 8 weeks old) that had been fed for two weeks during the dark-phase. Samples were collected at ZT0 (12h fed). This cohort was composed of 3 pairs of *RORc(t)$^{Cre}$ Vipr2$^{+/+}$* and *RORc(t)$^{Cre}$ Vipr2$^{fl/fl}$* littermates, housed in 3 different cages. **c,** Phylogenetic profile of the microbiota composition in the ileal fecal material of *RORc(t)$^{Cre}$ Vipr22$^{+/+}$* or *RORc(t)$^{Cre}$ Vipr2$^{fl/fl}$* mice after 12h of feeding. **d,** Weighted Unifrac Principal coordinate analysis (PCoA) of 16S rRNA composition in fecal pellet of the same mice described above from (*RORc(t)$^{Cre}$ Vipr2$^{+/+}$* or *RORc(t)$^{Cre}$ Vipr2$^{fl/fl}$* mice). **e,** Phylogenetic profile of the microbiota composition in the fecal pellet of *RORc(t)$^{Cre}$ Vipr2$^{+/+}$* or *RORc(t)$^{Cre}$ Vipr2$^{fl/fl}$* mice after 12h of feeding.

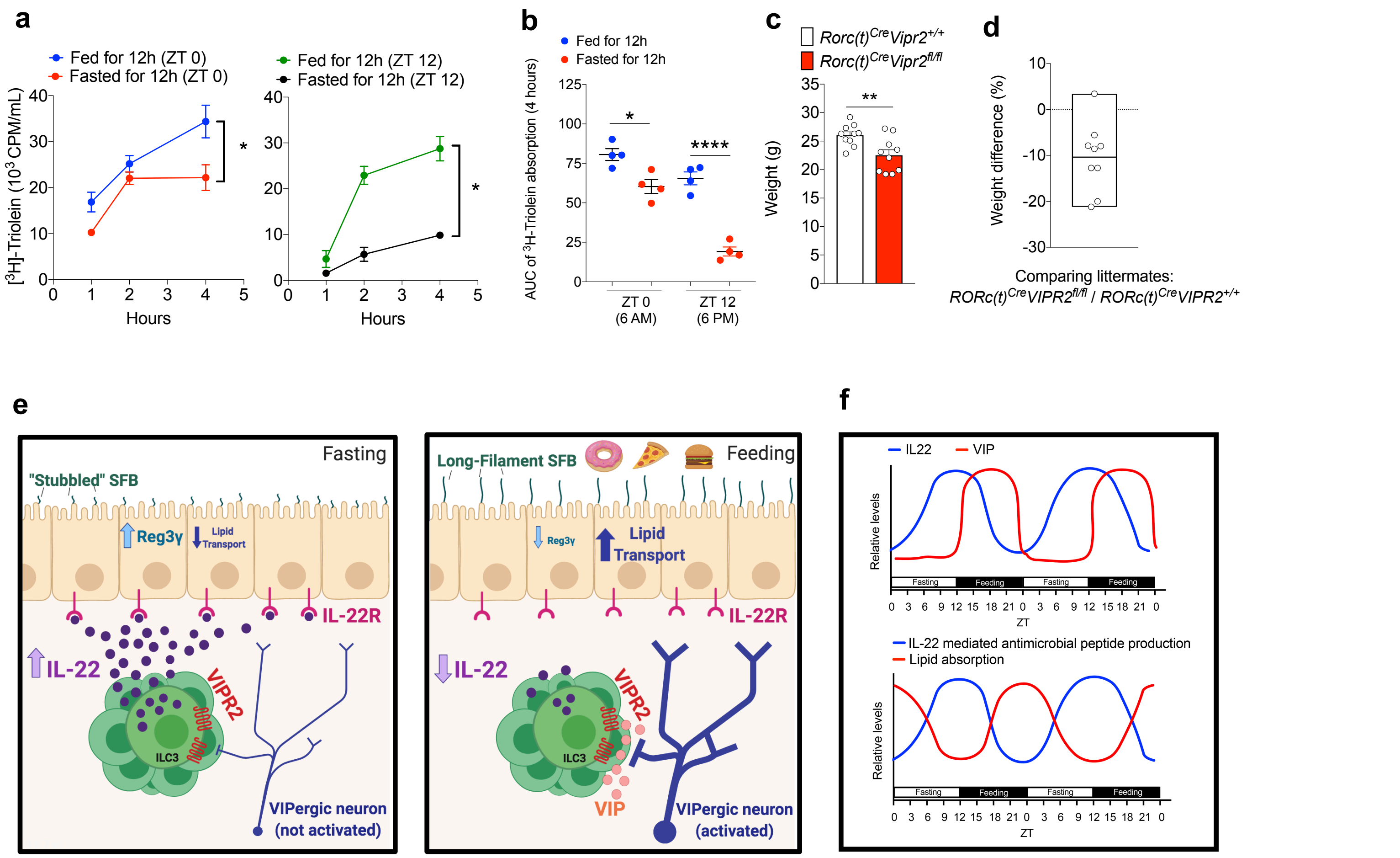


**Extended Data Figure 11. Feeding increase the efficiency of triglyceride absorption. a, b,** Plasma $^3$H CPM (counts per minute) in mice fed or fasted for 12h during the light-phase (ZT 0 – ZT 12, red and green circles) or during the dark-phase (ZT 12 – ZT 0, blue and black circles) and then gavaged with $^3$H-triolein and sampled at different times (**a**); and AUC during 4h **(b)**. AUC: Area under the curve per mL of plasma. N=4 mice per group, representative of two independent experiments, **P* < 0.05 and *****P* < 0.001 (two-way ANOVA). **c,** Weight of 10 littermate/cagemate pairs of *RORc(t)$^{Cre}$ Vipr2$^{+/+}$* and *RORc(t)$^{Cre}$ Vipr2$^{fl/fl}$* mice under regular chow diet (male, 9 weeks old). Representative of two independent experiments, ***P* < 0.01 (*t-test*). **d,** Weight difference (%) between each *RORc(t)$^{Cre}$ Vipr2$^{fl/fl}$* mouse compared to its paired *RORc(t)$^{Cre}$ Vipr2$^{+/+}$* littermate/cagemate. **e,f,** Graphical abstract **(e)** and theoretical model **(f)** of the observations described created with BioRender.
